# Community-level signatures of ecological succession in natural bacterial communities

**DOI:** 10.1101/636233

**Authors:** Alberto Pascual-García, Thomas Bell

## Abstract

A central goal in microbial ecology is to simplify the extraordinary biodiversity that inhabits natural environments into ecologically coherent units. We present an integrative top-down analysis of over 700 bacterial communities sampled from water-filled beech tree-holes in the United Kingdom at distances between <5m to >100km, combining an analyses of community composition (16S rRNA sequencing) with assays of community functional capacity (exo-enzymatic activities, ATP production, CO2 dissipation and yield). The communities were grown in laboratory conditions in a complex growth medium, what allowed us to investigate the relationship between composition and function, excluding confounding environmental factors. We found a distance-decay trend in the similarity of the communities, and simulated data allowed us to reject the hypothesis that stochastic processes dominated the assembly of the communities, suggesting that niche effects prevailed. Consistent with this idea, clustering of communities lead us to identify six distinct community classes encompassing samples collected at often distant locations. Using structural equation modelling, we explored how functions were interrelated, demonstrating that a representative functional signature can be associated with each community class. We obtained a more mechanistic understanding of the classes using metagenomic predictions with PiCRUST. Notably, this approach allowed us to show that these classes contain distinct genetic repertoires reflecting community-level ecological strategies. We finally formulated an over-arching ecological hypothesis about how local conditions drive succession in these habitats. The ecological strategies resemble the classical distinction between r- and K-strategists and could be extrapolated to other systems, suggesting that a coarse grained picture of microbial ecological succession may be explained by relatively simple ecological mechanisms.

## Introduction

The microbial communities inhabiting natural environments are unmanageably complex. It is therefore difficult to establish causal relationships between community composition, environmental conditions and ecosystem functions (such as rates of biogeochemical cycles) because of the large number of factors influencing these relationships. There is great interest in developing methods that reduce this complexity in order to understand whether there are predictable changes in community composition across space and time, and whether those differences alter microbe-associated ecosystem functioning. The most common approach has been to search for physical (e.g. disturbance) and chemical (e.g. pH) features that correlate with community structure and function. This approach has often been successful in identifying some major major differences among bacterial communities associated with different habitats [1] and some of the edaphic correlates [2]. However, even if significant correlations between environmental variables and microbial functioning are found, we are still far from understanding the underlying biological mechanisms explaining these relationships. For instance, adding variables such as biomass or diversity to models in which environmental variables are good predictors of function do not strongly improve model predictions [3], suggesting that there is a need for variables that increase the accuracy of biological processes [4].

The development of a more mechanistic picture is hindered for several reasons, such as difficulties in identifying the relative role of stochastic and deterministic processes in shaping microbial communities [5, 6, 7], the pervasiveness of functional redundancy [8, 9], and of priority effects [10]. In addition, it is often difficult to identify which functions to assess. Microbes inhabitating a host sometimes have a substantial impact on host performance, for example turning a “healthy” into a “diseased” host [11]. Such extreme impacts of individual taxa make it relatively simple to infer a direct link between community composition and function. In open, natural environments (e.g. soil, lakes, oceans), the impact of individual taxa on ecosystem functions is often minor, and generalisations may depend on subjective choices of which functions to measure.

An important step forward comes from manipulative experiments in natural environments, that have identified variables such as pH [12], salinity [8], sources of energy [13], the number of species [14], and environmental complexity [4] as key players in the relationship between bacterial community structure and functioning. Improved control can be obtained by “domesticating” communities surveyed from natural environments by growing them in a synthetic (albeit complex) environment, and quantifying their functioning under such controlled conditions [15, 16]. With these experiments it becomes possible to directly test the hypothesis that more similar communities have more similar functions without the confounding influence of extrinsic environmental conditions.

Community similarity can be assessed using a rich array of analytic tools that identify *β*–diversity clusters within multivariate datasets, such as the detection of communities in species co-occurrences networks [17] or the reduction of the dimensionality of *β*–diversity similarities [18]. These approaches have been pervasive in the medical microbiome literature, for example in the search for “enterotypes”-i.e. whether individuals are characterised by diagnostic sets of species representing alternative community states [18, 19] which, in this paper, we call “classes” of communities. The existence of classes in communities sampled from different locations may be due to variable environmental conditions that select for different taxa, or may be explained more parsimoniously by stochastic processes together with strong dispersal limitation [20]. Deciphering the likelihood of different ecological mechanisms can be assessed by adopting a suitable null model, see e.g. [21]. Community classes arising from environmental selection would also be functionally different, whereas we would not expect functioning to differ among community classes created by stochastic processes.

Once classes and functional differences have been identified, it is possible to step down into key biological processes focusing on the genetic repertoires of the constituent taxa [22]. Investigating the dominant genes present in the different community classes allows explanations of functional differences and the determination of ecological strategies. For example, community classes that differ in genes related with environmental sensing, degradation of extracellular substrates, or metabolic preferences, could be used as hypotheses of the molecular mechanisms responsible for functional differences. Therefore, the last step aims to explain how the functional and genetic differences arise from the prevailing environmental conditions [23], and could point to the specific environmental parameters that could be measured. This approach solves the problem of measuring many environmental parameters in the hope that some will be significantly associated with community structure or ecosystem functioning. Lack of any clear functional differentiation among community classes is also informative, and would indicate alternative community states with redundant functions [24, 25, 26]. Such redundancy could arise in the absence of environmental variability, which could also help explain the lack of a dominant environmental axis that explains variation in composition.

In this work we followed the above pipeline investigating a large dataset consisting of more than 700 samples of rainwater-filled puddles (phytotelmata) that can form at the base of beech trees. The bacterial communities present in the tree-holes are key players in the decomposition of leaf litter, and therefore of great interests more broadly for understanding decomposition in forest soils and riparian zones. This is an ideal system to follow this procedure given the relatively similar conditions found across different locations, making it unique in terms of replicability of a natural aquatic environment [27, 28], and its relatively low diversity. Indeed, while effects caused by environmental variation on phytotelmata ecosystems has been investigated in meio- and macrofaunal communities [28], the influence in microbial communities is largely unknown. Moreover, emphasis was made for understanding bottom-up drivers of tree-holes diversity like nutrients [29, 30, 31], but top-down approaches that may help us to understand other drivers of microbial composition like stochastic dispersion or interactions have received less attention [28].

Previous work using this dataset showed that larger (smaller) taxa abundances influenced broad (narrow) community-level functions [32]. Here we aim to illuminate the mechanistic basis of this relationship. The large dataset allowed us to study natural variation in bacterial community composition through the top-down categorisation of communities into classes. We then linked the classes with bacterial functioning, analysing a set of community-level functional profiles obtained from laboratory assays of the same communities [32], and investigating whether the classes differed in their functional capacity. Instead of focusing on each function individually, we investigated how the functional profiles varied across the community classes. We then used metagenomic information to understand whether similar compositions and functions were translated into different classes of genetic repertoires.

We found significant differences in the genetic repertoires and functional measurements among classes, that we interpreted in the context of changing environmental conditions. In deciduous forests, bacterial life on the forest floor is characterised by seasonal and daily changes to temperature and resource availability. We therefore address whether differences in the communities are due to the historical processes at the different geographic locations, or if they are rather more influenced by contingent local conditions. These factors are often difficult to resolve [26] but may both be important due to the high temporal variability in these systems, as observed in compost ecosystems [33]. Interestingly, interpreting the signatures found in the functional measurements and in the genetic repertoires led us to hypothesize the existence of community-level ecological strategies reflecting an ecological succession driven by local environmental dynamics of the tree-holes. These ecological strategies ressemble the classical distinction between r– and K–strategists described for single species [34].

## Results

### Microbial communities classes are determined by local conditions

We analysed 753 bacterial communities sampled from water-filled beech tree-holes in the South West of UK [32] (see Suppl. Table 1). Communities were grown in a medium made of beech leaves as substrate for seven days and then their composition interrogated through 16S rRNA sequencing (see Methods). We analysed the *β*–diversity of these communities according to two different metrics: the Jensen-Shannon divergence (*D*_JSD_) [35], and a transformation of the SparCC metric (*D*_SparCC_, see Methods [36]). Spatial distances and the two *β*–diversity distances were significantly correlated (Mantel test: *r* = 0.21; *p* < 10^−3^ for D_SparCC_ and *r* = 0.19; *p* < 10^−3^ for *D*_JSD_). To test if this trend was maintained across the different scales, we clustered samples that were closer in space, and retrieved the classifications found at 10 distance thresholds spanning 5 orders of magnitude (from <5 m to >100km). We used three statistics (ANOSIM, MRPP and PERMANOVA [37, 38], see Methods) to test whether the *β*–diversity distances within clusters are significantly smaller than those between clusters for the 10 classifications. In all cases, the three tests supported the hypothesis that communities within locations were significantly more similar than between locations (permutation tests, *p* < 10^−3^, see Suppl. Fig. 1). This spatial autocorrelation may be indicative of an important role for stochastic assemblage and dispersal limitation [20].

We studied how the different statistics changed across the 10 classifications. We observed an increase in the mean distances within clusters (quantified with the MRPP statistics) and a decay in the ANOSIM *R* statistics (Fig. 1 c and d), while PERMANOVA remains roughly constant across scales (Suppl. Fig. 2). To interpret these trends we analysed the behaviour of these metrics with synthetic data in which artificial *β*–diversity distances matrices were generated under different assumptions on the mean and variances within and between locations across scales (Suppl. Figs. 3—6). We found that the most informative observation is the ANOSIM-*R* decay when the variance of the simulated distances were large (Suppl. Figs. 6 middle column, bottom) a pattern consistent with the one observed for the real data (Fig. 1d). To give a sense of the implications of this finding, in Suppl. Fig. 7 we show that for such variance in the generated distances, approximately 3% of the *β*–diversity distances between samples located at more than 100 km. away should be as high as those found in communities samples within 5 m. of distance.

**Figure 1:**
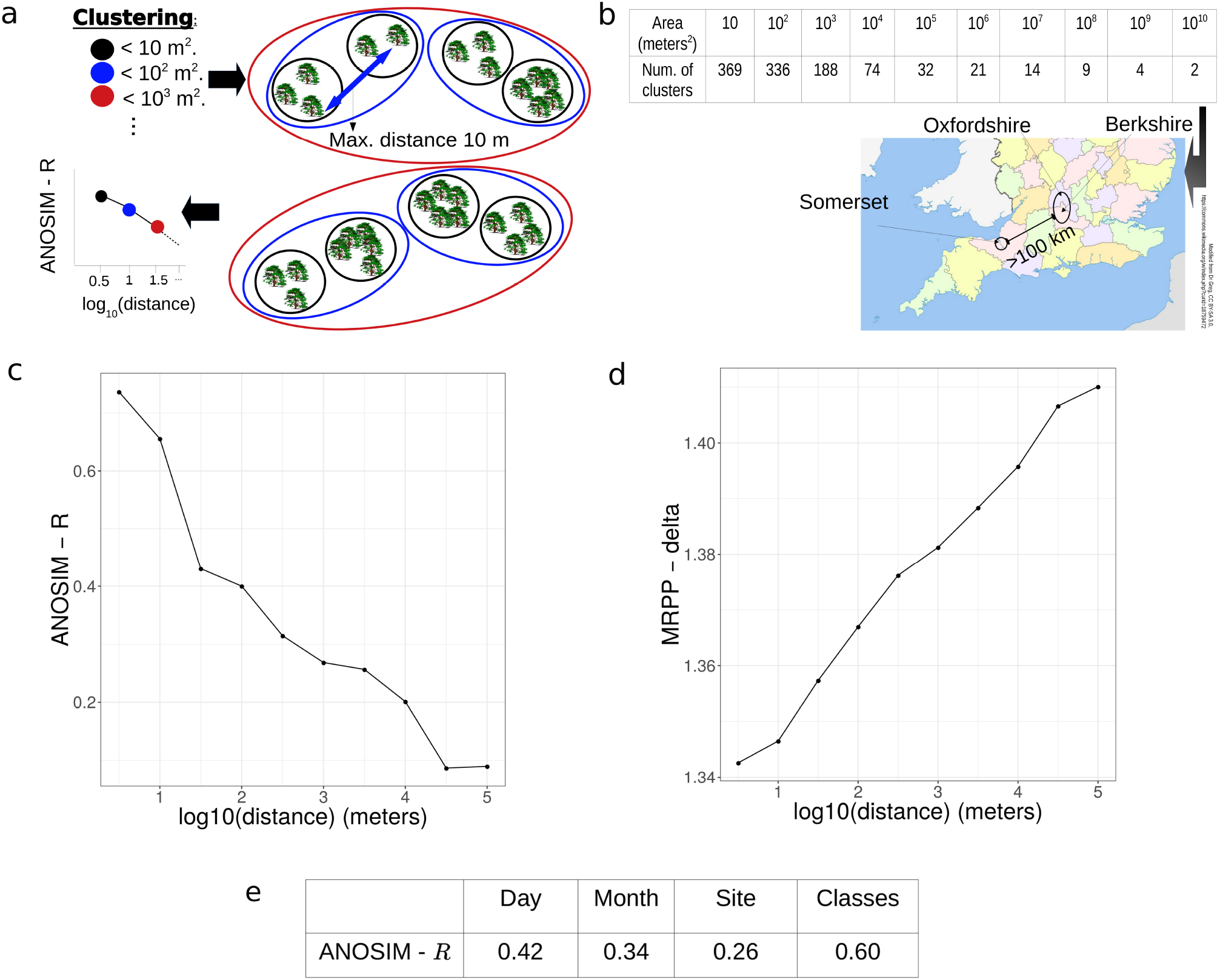
Analysis of decay of communities’ similarity with distance. (A) Illustration of the procedure. Trees are clustered within areas A of increasing sizes, leading to one classification every order of magnitude (ellipses of different colours). We approximate these areas, stopping the clustering at spatial distances thresholds equal to 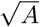, e.g. the classification for areas of 100 *m*^2^ (blue ellipses) is obtained stopping at *d* = 10*m*. For each classification, an ANOSIM and mrpp value was computed to test if the similarity of the communities within the clusters is larger than between clusters, and the statistics were plotted against the spatial threshold. (B) Number of clusters at each spatial threshold. The three counties from which the communities were sampled are shown in the map. The classification at the last threshold consists of two clusters separated by more than 100km. (C and D) Observed ANOSIM and mrpp statistics for each of the distance thresholds, reflecting a decay in the similarity of the communities (see Main Text for details). (E) ANOSIM values for communities clustered according to the day and month in which were sampled, the site from which they were collected (optimal classification of spatial distances), and the six community classes (optimal classification of *β*–diversity distances).

This result implies high levels of dispersal that are difficult to explain for tree-hole bacteria. We explored the alternative hypothesis: that similar communities found at distant locations result from similar underlying environmental conditions. We performed unsupervised clusterings with *D*_JSD_ and *D*_SparCC_, revealing in both cases six distinct community classes (Fig. 2A and B, Suppl. Fig. 8 and Suppl. Table 2 for global characteristic metrics such as diversity). The whole set of communities are dominated by *Proteobacteria*, and the community classes were distinguished at the genus level (95% sequence similarity), including a higher presence of the genus *Klebsiella* and *Pantoea* (classes 1, red; and class 3, pink); *Paenibacillus* and *Sphingobioum* (class 2, green); *Serratia* (class 5 blue); *Sphingomonas, Streptomyces* and *Pseudomonas* (classes 4, yellow) and low abundant genera like *Brevundimonas* and *Herbaspirillum* and, again, *Pseudomonas* for class 6 (grey). In the following, we refer to class 1 (red) as the reference class because it encompassed the largest number of communities (Suppl. Table 2). We refer to the remaining communities by their most distinctive taxon as *Paenibacillus* (class 2), *Klebsiella* (class 3), *Streptomyces* (class 4), *Serratia* (class 5). For class 6 we observed that, although the *Pseudomonas* genus was also high in other communities, classes 4 and 6 were dominated specifically by *Pseudomonas putida* (Suppl. Fig. 9), which we selected as representative of class 6. In Ref. [39] we use a network approach to identify modules of coocurring species that confirm the key role of the taxa selected as representatives.

**Figure 2:**
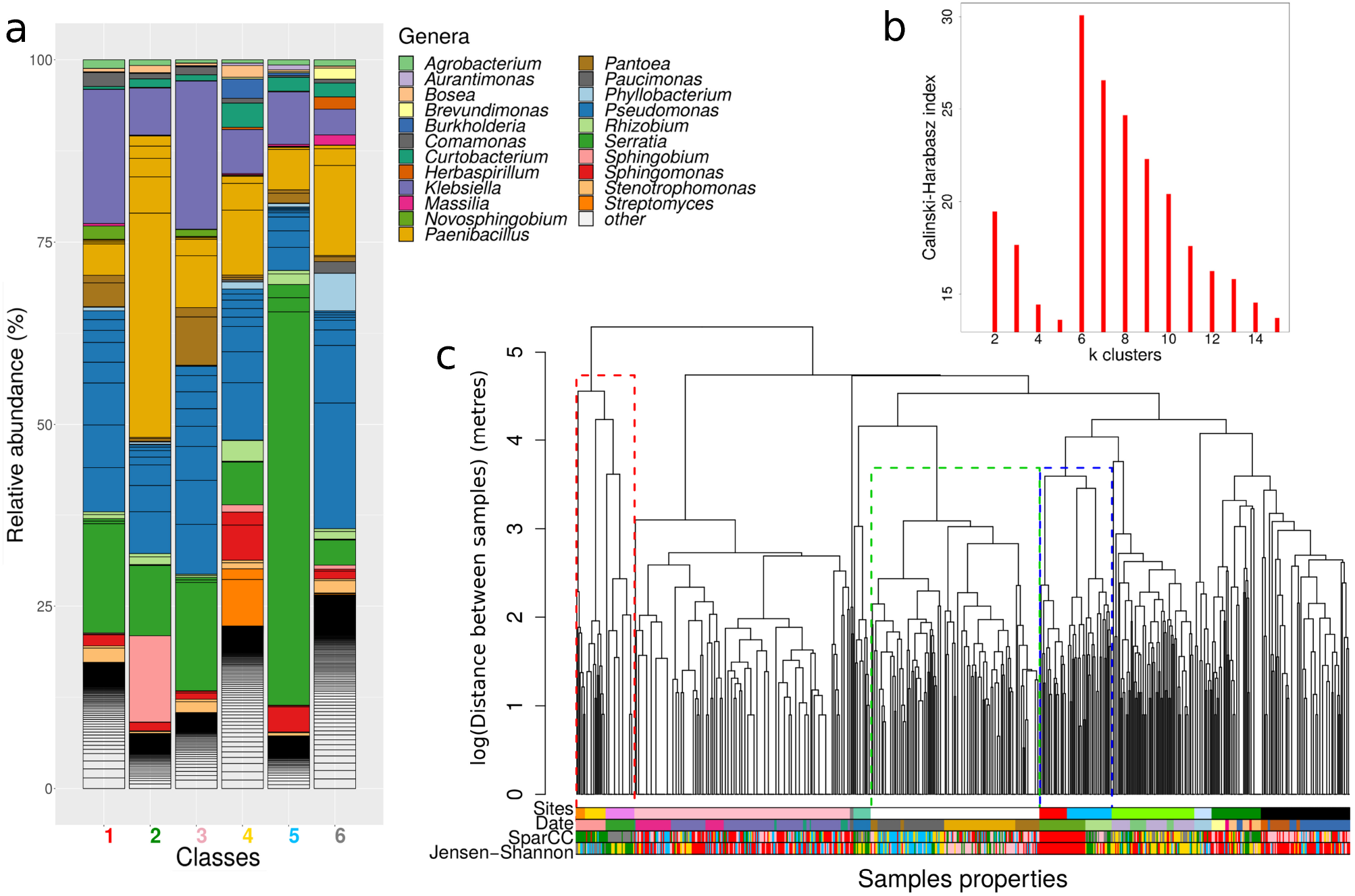
Communities classes description and relation with sampling sites. (A) Genus frequencies in each of the six community classes found after unsupervised clustering of the communities using *D*_SparCC_. Only the 15 most abundant genera in each community class are shown, with the remainder classified as Others. (B) Calisnki-Harabasz index versus the number of final clusters in the classification, showing a maximum at six classes. (C) The dendrogram represents the clustering of the samples (leafs in the tree) according to their spatial distances. The coloured bars represent membership of each sample to different classifications (from top to bottom): the sites, the day of sampling collection (Date), and the classes found using either *D*_SparCC_ and *D*_JSD_. Dotted rectangles in the dendrogram indicate examples of: (green) locations where one site was sampled on different days; (blue) two sites sampled on the same day: (red) a mix of both situations. It is apparent that the date is a better match to the community classes than the site, which is confirmed with a comparison between the classifications (Suppl. Table 3).

We illustrate how these classes are distributed in space by representing the class identity of each community in a coloured bar, alongside the Site and Date in which the community was sampled (Fig. 2C). As expected from the previous analysis, some communities belong to the same class even if they were distant in space (see examples in Suppl. Fig. 10). For the latter, in Fig. 2C we highlight with dotted rectangles some of the cases in which there is a better correspondence with the date of sampling than with the site. To test this observation, we showed that a classification based on the date of sampling is consistently more *β*–similar to the diversity classes than the site (Suppl. Table. 3). Moreover, computing the ANOSIM statistics when tree-holes are clustered according to sampling location (Site values in Fig. 2e) or according to the sampling date (Day and Month values) consistently show that the specific date (Day) is more informative than the site. The Day was also more informative than the Month, suggesting that seasonal environmental conditions were not the main drivers of the similarities, but were rather due to daily variation in local conditions. Notably, the value we obtain for the ANOSIM statistics when the classification considered are the community classes found, reaches the same value than the one found at 50 m (Fig. 1c). In summary, the classification successfully grouped communities into just six groups, with communties often separated by far more than 50 m. In addition, the date of sampling is more informative than the sites. Taken together, the results suggest that the classes capture similarities in local environmental conditions even in tree-holes that were spatially separated by considerable distances.

### Community classes reflect different functional performances

If environmental conditions determine compositional differences in the communities, we expect that these differences are translated into different community functional capacities, a question that we investigate analysing data that quantify the function performance of these communities [32]. The sampled communities were cryo-preserved after sequencing, and later revived in a medium made of beech leaves as substrate. Cells were grown for seven days while monitoring CO2 dissipation and, after this period the following measurements were taken: community cell counts, community metabolic capacity (measured as ATP concentration), and community capacity to secrete four ecologically relevant exoenzymes [40] related with i) uptake of carbon: xylosidase (X) and *β*–glucosidase (G); ii) carbon and nitrogen: *β*–chitinase (N); and iii) phosphate: phosphatase (P).

Visual inspection indicated substantial differences in the functional capacities among the community classes. In some cases, communities belonging to different classes clearly separated, as shown in the histograms in Suppl. Fig. 12. Therefore, we explored if these differences among the community classes were significant using structural equation models (SEM) [41]. A model that made no distinction among classes yielded an excellent fit (RMSEA<10^−3^, CI=[0-0.023], AIC=7493 see Methods and Suppl. Fig. 14). From this structural model we next considered models with different parameters for each community class (up to six parameters per pathway, see Methods). We then investigated which model better explained the data according to 3 scenarios: i) a model in which all the parameters were constrained to be the same for all the classes; ii) a model in which each class had a different parameter for each pathway; iii) an intermediate model, in which some parameters were constrained for some classes. Accounting for penalizations for models with more degrees of freedom (RMSEA< 10^−3^, CI=[0-0.035], AIC=6658, see Methods and Suppl. Material), the best model belonged to the scenario (iii) (see Fig. 3 and Suppl. Table 6). This result supports the hypothesis that the classes had differentiated functional capacities (Suppl. Table 7 and Suppl. Fig. 15).

**Figure 3:**
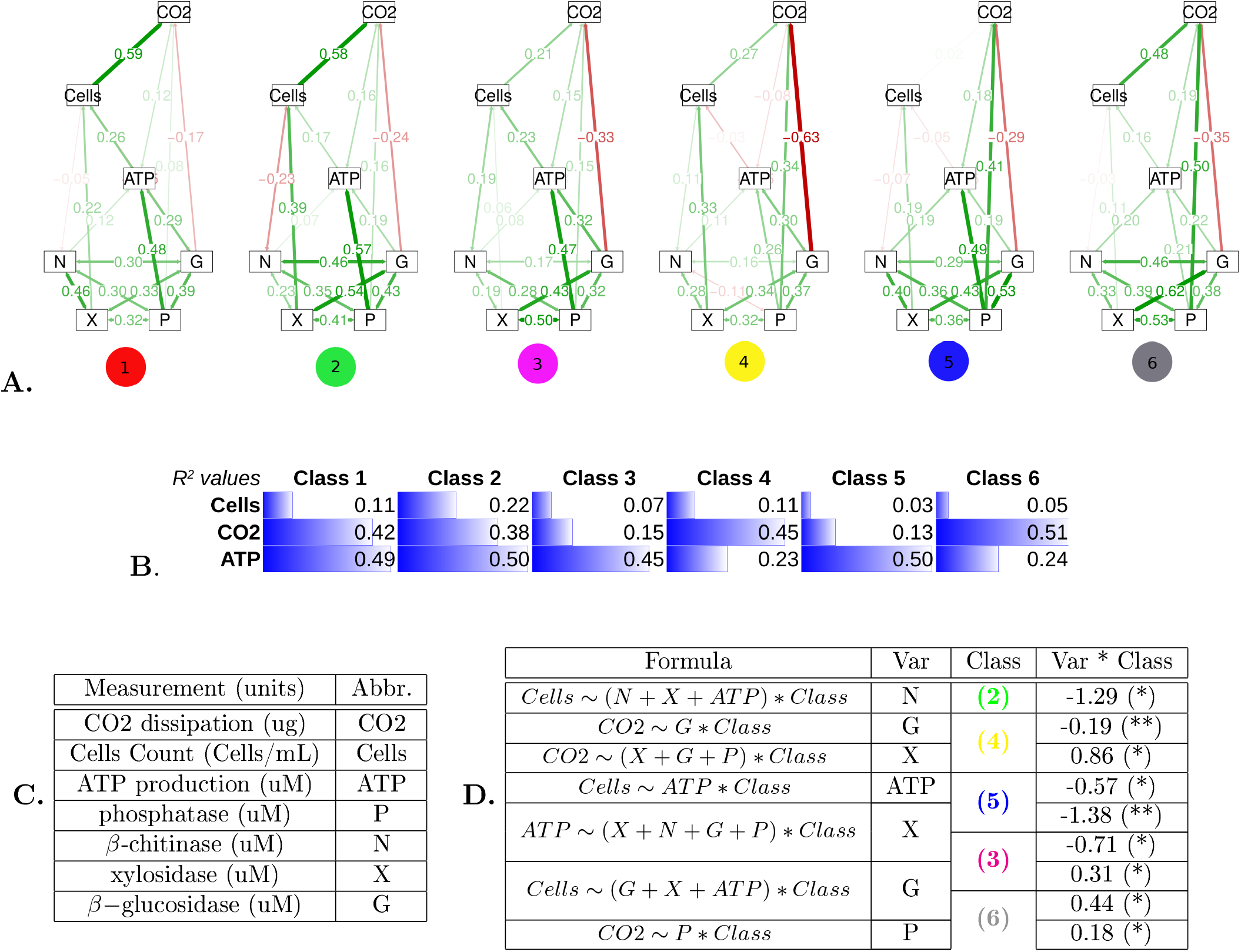
Functional capacity of the the communities classes. (A) Diagrams of the final SEMs. The standarized coefficients for each of the six classes are shown on the corresponding pathway. (B) *R*^2^ values for the three endogenous variables in the SEM model. (C) Experimental measurements used in this study and their abbreviations. (D) Causal analysis of the influence of the exogenous variable (Var) for each SEM pathway (determined by the response variable and Var in Formula) when the identity of the classes is included as a factor. Confounding factors involved in the pathway are also included (remainder variables in Formula). The column Class reflects the class identity of any significant coefficient found with its value shown in the column Var * Class (significance code: * = p-val<0.01, ** = pval<0.001).

The model showed that measurements related with uptake of nutrients were all exogenous, including ATP production, cell yields and CO2 production (Fig. 3). In addition, ATP production influenced yield, which in turn influenced CO2. Among exoenzyme variables, N influenced ATP and, notably, only X affected yield, while G and P influenced both ATP and CO2. Analysis and clustering of the standarized partial regression coefficients (Suppl. Fig. 11) established a classification of the classes in three groups according to their function, with *Paenibacillus* and the reference classes being the most similar, then *Streptomyces* and *P. putida* classes, and *Klebsiella* and *Serratia*.

We then explored if distinctive pathways for each class could be determined. Given the complexity of the SEM models, we first ruled out the possibility that differences in pathway coefficients were due to the influence of other (confounding) variables. To control for this possibility, for each pair of endogenous-exogenous variables, we searched for its set of confounding variables with dagitty [42]. Next, for each pair of variables involved in a pathway, we performed a linear regression including its adjustment set of confounding factors, and an interaction term with a factor coding for the different classes. Coefficients should be interpreted as deviations with respect to the reference class (see Methods). The significant interaction terms (Fig. 3, panel D) show how the relationships among the functional variables differed among the community classes. For example, the analysis revealed that cell yield was negatively influenced by *β*–chitinase activity for the *Paenibacillus* class, for ATP production for the *Serratia* class, while being positive related with *β*–glucosidase for the classes of *Klebsiella* and *P. putida*. We therefore concluded that the community classes had significantly different functional capacities, thus producing different relationships in the models.

### Community classes depict different genetic repertoires

To get a more mechanistic understanding of the above results, we analysed the genetic repertoire of each community class by performing metagenomic predictions with PiCRUST [43], and further statistical analysis with STAMP [44]. The Nearest Sequence Taxon Index is 0.059, reflecting a high quality prediction [43]. This is because most of the dominant genera in this system are found in gut microbiomes (e.g. Fig. 5 in Ref. [45]).

The fraction of exo-enzymatic genes belonging to *Paenibacillus, Streptomyces* and *P. putida* classes was significantly larger than the fractions found for the *Klebsiella, Serratia* classes and the reference class, suggesting that the former classes are specialized in degrading a wider array of substrates (Fig. 4).

**Figure 4:**
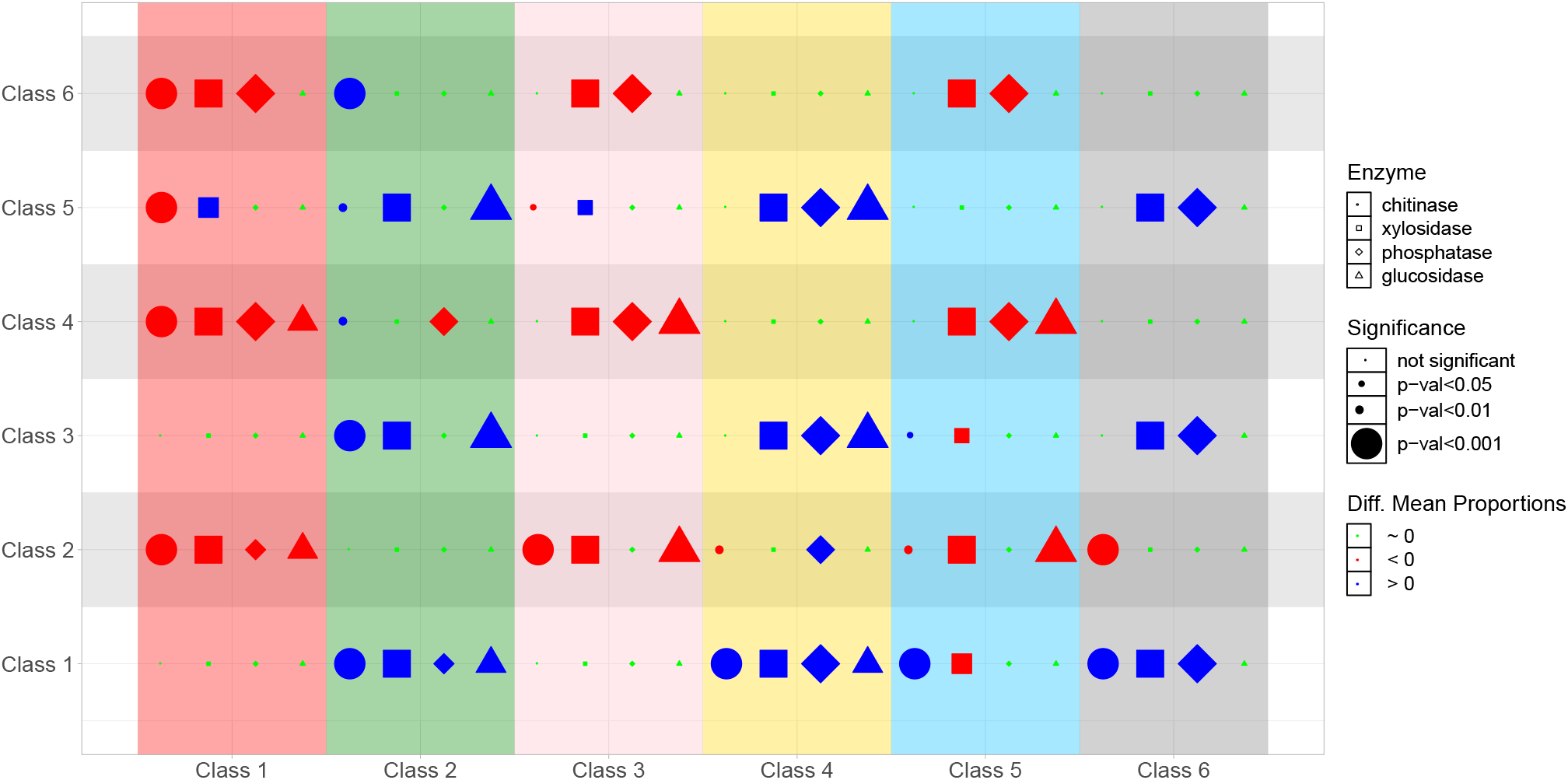
Summary of post-hoc analysis of exo-enzymatic genes predictions. The significance of differerences in mean proportions of chitinase (circles), *β*-xylosidase (squares), *β*-glucosidase (triangles) and phosphatase (dianionds) were tested across all pairwise combinations of community classes. Red (blue) symbols denote that the class in the correspondent column has a significantly lower (higher) mean proportion of the enzyme for the class shown in the row. The size of the symbol is larger for more significant differences. All tests are provided in Suppl. Fig. 4.

Clustering the KO annotations into KEGG pathways (see Methods) showed that the 6 community classes differed in their genetic repertoires. Furthermore, these divergent genetic repertoires suggested different ecological adaptations, which are summarized in Fig. 5. Consistent with PCA analysis of the KEGG pathways (Suppl. Figs. 18-21), we divided the classes in two groups: the reference, *Klebsiella* and *Serratia* classes carried the genetic machinery for fast growth, while *Paenibacillus, Streptomyces* and *P. putida* classes carried the genetic machinery for autonomous amino-acid biosynthesis. Evidence for fast growth in the reference, *Klebsiella* and *Serratia* classes comes from the large fraction of genes related with genetic information processing (Suppl. Fig. 24), mostly related with DNA replication such as DNA replication proteins genes, transcription factors, mismatch repair, homologous recombination genes or ribosome biogenesis—the latter being a good genetic predictor of fast growth [46]. Secondly, communities from these classes also carried a larger fraction of genes related with intake of readily-available extracellular compounds (Suppl. Fig. 25), including ABC transporters, phosphotransferase system, or peptidases, and environmental adaptations including motility proteins, synthesis of siderophores, and the two-component systems. Rapid replication often requires a more accurate control of protein folding and trafficking. Consistent with this hypothesis, we found a significantly inflated fraction of genes involved in folding stability, sorting and degradation, including chaperones and genes involved in the phosphorelay system (Suppl. Fig. 26).

**Figure 5:**
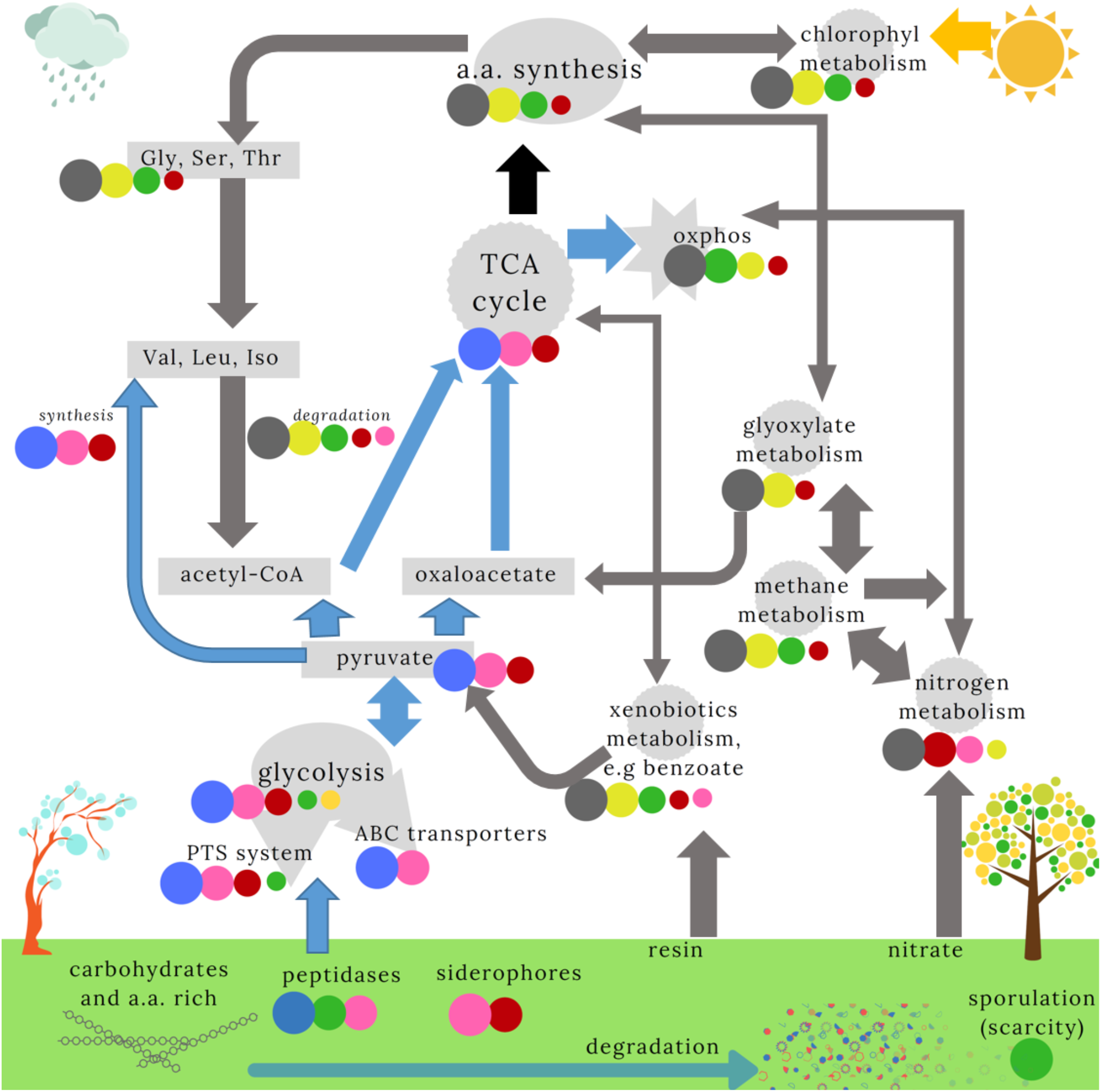
Scheme describing the genetic repertoires of the six community classes. Pathways from the KEGG database that were most relevant for describing the classes are shown. We ranked the mean proportion of genes in each pathway (see Suppl. Material), indicated by the size of the circles (Fig. 4). Only classes with significant pairwise post-hoc comparisons are shown. *P. putida* (gray) and *Serratia* (blue) classes appear to dominate orthogonal pathways. We therefore indicated how the pathways were influenced by the dominant community (indicated by the arrow colour). The link between the TCA cycle and amino acid synthesis (black arrow) is unclear. We further illustrate the substrates and hypothetical environmental conditions expected for *Serratia* (rain/cold) and *P. putida* (dry/hot) communities. We suggest the other communities are intermediates between these two classes. The *Paenibacillus* class, with a large number of sporulation and germination genes, may reflect conditions of very low nutrient availability.

A second series of evidences pointing towards orthogonal ecological strategies came from differences in the metabolic pathways associated with the community classes. *Serratia*-dominated class (5) had an inflated fraction of genes related to carbohydrate degradation, including genes involved in glycolysis and in the trycaborxylic acid (TCA) cycle (Suppl. Fig. 27). In contrast, the *Paenibacillus, Streptomyces* and *P. putida* classes were associated with genes involved in alternative pathways like nitrogen/methane metabolism, and in secondary metabolic pathways related with degradation of xenobiotics/chlorophyl metabolism. Notably, the genes involved in the exoenzymes that were experimentally assayed were higher in these classes, suggesting that they were adapted to environments with more recalcitrant nutrients (Fig. 5). In addition, *Paenibacillus, Streptomyces* and *P. putida* classes had a remarkable repertoire of genes for amino acids biosynthesis–possibly at odds with the reference class and *Klebsiella*, and *Serratia* classes, which invested in proteases for amino acid uptake (Suppl. Fig. 28 and 29). The apparently low glycolytic capabilities of these communities could result in pyruvate deficiencies, which would hindered the production of sufficient acetyl-CoA and oxaloacctatc required to activate the TCA cycle. Consistent with this observation, we observed that these communities exhibited a significantly larger proportion of genes related with glyoxylate metabolism and degradation of benzoate, that may be used as alternatives to glycolysis (Suppl. Fig. 30). Finally, we observed that communities in the reference class and Klebsiella and *Serratia* classes had a significantly larger repertoire of genes needed to synthesise amino acids requiring pyruvate (valine, leucine and isoleucine), and which, according to our interpretation, they would generate through glycolysis (Suppl. Fig. 27). By contrast. *Paenibacillus, Streptomyces* and *P. putida* classes had a significantly larger proportion of genes used to degrade these amino acids (Suppl. Fig. 28) and hence, either they take these essential amino acids from the environment or they generate them from other pathways. Consistent with this observation, genes in these classes were enriched for glycine, serine and threonine metabolism (Suppl. Fig. 28). through which it is possible to obtain valine, leucine and isoleucine, and which could provide an alternative source of acetyl-CoA (Fig. 5).

## Discussion

Our analysis of a large set of tree-hole bacterial communities found a strong distance-decay in the similarity of the communities across several orders of magnitude. The existence of spatial autocorrelation has previously reported in soil environments [47, 12], but this study extends the findings to scales above the short distances (<10m) previously reported [47]. The high ANOSIM statistics require unrealistic levels of dispersal for the pattern to be explained by stochastic processes alone (Suppl. Figs. 28), and therefore points towards a hypothesis that similar environmental conditions occur at distant locations.

We observed that the communities could be arranged into classes, and that the classes corresponded to the site and the date of collection, which are tightly entangled. There was a geographic influence that was particularly apparent at larger distances, which suggests the influence of broad environmental conditions and perhaps historical processes [26]. The finding is consistent with the idea that environmental conditions on a particular day strongly influenced species composition, consistent with previous findings on macro-invertebrate tree-holes communities [48]. Moreover, particular classes were found in different seasons suggesting that factors like temperature were of secondary importance, despite results highlighting their importance in similar systems [25]. Communities collected within the same day were more similar if they were closer in space, demonstrating significant spatial autocorrelation, which has previously been reported in soil environments [47, 12].

Laboratory experiments confirmed that these classes were associated with different functional capacities, which we believe strongly implies that the classes are ecologically meaningful subgroups. The result was compatible with a scenario of ecological succession in which there was a transition from communities dominated by r-strategists to K-strategists [49]. We suggest that early successional stages were characterized by the *Serratia* class. This class had a negative relationship between ATP and cell yield, indicating low resource use efficiency. In addition, investing in xylosidase had a much lower transfer into ATP production than for the reference class, implying a preference for labile substrates like sugar monomers. Analysis of the metagenome revealed pathways responsible for extracellular degradation and uptake of nutrients, and metabolic processes associated with glycolysis. The class also had many genes associated with environmental processing, fast replication and accurate molecular control of protein folding and trafficking. The mean Shannon diversity of communities belonging to this class was almost the lowest (Suppl. Table 2) which might be expected in a rich environment dominated by a few well adapted fast growers, consistent with the notion of r-strategists.

The next communities in the succession were the reference and *Klebsiella* classes. Although still sharing some of the features of the *Serratia* class, they had distinctive features such as a higher conversion of ATP into yield. Later successional stages were characterized by the *P. putida* and *Streptomyces* classes, exhibiting high respiration values. These classes contained an inflated fraction of genes related to oxidative phosphorylation and were able to synthesise most amino acids. They were also associated with secondary metabolic pathways that may be valuable in environments in which resources are low but where it is possible to scavenge the metabolic by-products of former inhabitants. This is particularly apparent for the *P. putida* class, which also had a higher Shannon diversity including many rare species, consistent with communities dominated by K-strategists competing for rare and heterogeneous resources.

Finally, the *Paenibacillus* class contained many of the metagenomic characteristics of the *P. putida* and *Streptomyces* classes. It was the class with lowest Shannon diversity, and also a large fraction of sporulation and germination genes (Suppl. Fig. 31). These results imply that these communities lived particularly unproductive environments. The laboratory results are consistent with this hypothesis: this community had the largest conversion of chitinase activity into yield, which may reflect its ability to take advantage of the remaining nutrients such as dead arthropod exoskeletons or fungi. Water volume is among the main driver of fungi sporulation in this system [50], which would match our interpretation. Taken together, the results imply this class is the last stage of the succession, where nutrients have been depleted to low levels.

There are several environmental conditions that might be driving succession. First, succession may be due to nutrients dynamics in the tree-holes. A main source of carbon is beech leaf litter, supports meio- and macrofaunal communities [51, 52]. Degradation of leaf litter would be compatible with the succession described. Following leaf fall, any imple sugars would rapidly be used over days to weeks, while starch and cellulose degrade much more slowly [53]. If this is the main driver of succession in tree-holes, we would expect a strong seasonal signal, with a class dominating in autumn. Our data do not support these observations because the month of the year was a relatively poor classifier of the samples, and members of the classes we identified were often from different times of year.

Second, succession may be due to patterns of rainfall. Rainwater can bring nitrogen, sulfate, and other ions into the tree-hole, but the pathway followed by the water (stemflow or throughfall) will influence the final chemical compositions [29, 31]. For example, flushing after heavy rain can reduce phosphate levels to a minimum [30], and labile orthophosphate is expected to increase at later successional stages [31]. In addition, a progressive acidification in tree-holes that do not receive water inputs for long periods is also expected due to nitrification [29, 31]. Rain pulses can therefore have rapid impacts on tree-hole conditions and may explain the similarity of some samples collected at the same date even at distant locations, while other properties of the tree-holes like size, litter content, and the modes of water collection may preclude complete synchronization.

We envisage a scenario in which rain events were the primary drivers of bacterial composition, illustrated in bottom-left corner of Fig. 5, which would be modified by tree-hole features (e.g. volume, leaf inputs). Rain would generate pulsed resources of different type and frequency [54], and tree-holes features would determine the rate of resource attenuation [55]. For instance, large tree-holes or those with large leaf contents would have a slower rate of succession since resources are depleted less rapidly. This hypothesis would explain why, on some dates, all the tree-holes had similar compositions (recent rain or long standing drought conditions) while, beyond that, the classes are distributed across different dates and sites (due to the differential tempos of succession in tree-holes with different features).

Dissolved oxygen may be a third environmental component that influences community composition. The *P. Putida* class were associated with genes involved in aerobic respiration and high levels of phosphate. We observed an increase in abundances of strict aerobes, including *Brevundimonas, Paucimonas* and *Phyllobacterium*. There was also an increase in genes related with metabolism of nitrate, methane, degradation of benzoate (likely associated with the presence of resines), or chlorophyl (which indicates an increase in photo-heterotrophs). This class might also be able to run the TCA cycle generating acetyl-CoA from acetate, and from the degradation of valine, leucine and isoleucine, further complemented with glyoxylate metabolism and the degradation of benzoate to generate oxaloacetate. Finally, the class was found in summer and winter, and clustered in specific areas. This makes it less likely that temperature is an important variable, and points towards the amount of water and oxygen as key variables. This observation could also hold for the *Paenibacillus* class, for which long drought periods could lead to lack of water regardless of other tree-holes features (Suppl. Fig. 10).

We cannot rule out other site-based conditions like the type of forest management. A study analysing this factor did not find substantial differences in enzymatic activities despite different community compositions [56], perhaps because the low number of samples did not bring sufficient resolution. Another possible local influence for the composition are trophic ecological interactions, like the prevalence of invertebrates in certain areas (e.g. mosquito larvae) [57]. Insects with flying stages may also influence dispersal among tree-holes, which might contribute to microbial community similarity within a site [58], resulting in a metacommunity structure [48].

The approach taken here provides detailed insights into the community ecology of the bacterial communities inhabiting rainwater pools. By identifying community classes a priori, we were able to piece together the natural history of this environment from the perspective of the bacterial taxa. The spatial and temporal distribution of these classes, combined with the inferred metagenomes, indicate how environmental conditions reflect the metabolic specialisations of the dominant members. In this way, we were able to identify classes resembling r-vs. K-strategists [49] inhabiting tree-holes that were at different successional stages, a distinction also apparent in gut’s microbiomes [?]. Although this is no doubt an oversimplification, in general we find this conceptual framework is useful for microbes [34], since this ecological dichotomy may well be supported by thermodynamic [59] and protein-allocation trade-offs [60], which might also underlie other observed life history tradeoffs in microbes (e.g. olitgotrophic vs. copiotrophic strategies, [61]). We believe this approach therefore holds great promise for reducing the complexity of microbial community datasets [62], particularly in systems where the microbial communities have not yet been well characterised. In these systems, the approach we have used would generate hypotheses that could become the focus of future experiments or more detailed sampling strategies, therefore forming the basis of a bottom-up synthetic ecology that can be predictive in the wild.

## Methods

### Dataset

We analyzed 753 bacterial communities sampled in from rainwater-filled beech tree-holes (Fagus spp.) from different locations in the South West of United Kingdom, see Suppl. Table 1. 95% of the samples were collected between 28 of August and 03 of December 2013, being the remaining 5% collected in April 2014. Spatial distances between samples spanned five orders of magnitude (from <1m to > 100km). Sampled communities were grown in standard laboratory conditions using a tea of beech leaves as a substrate. After 7 days of growth, communities were characterized by sequencing 16S rRNA amplicon libraries from Ref. [32]. We considered only samples with more than 10K reads, and species with fewer than 100 reads across all samples were removed. This led to a final dataset comprising 680 samples and 618 Operational Taxonomic Units (OTUs) at the 97% of 16 rRNA sequence similarity. In previous work [32], four replicates of each of these communities were revived and regrown in the same media, further supplemented with low quantities of 4 substrates labelled with 4-methylumbelliferon. After 7 days, the experiments quantified the capacity of the communities to degrade xylosidase (abreviated X in the text, cleaves the labile substrate xylose, a monomer prevalent in hemicellulose), of *β*-chitinase (N, breaks down chitin, which is the major component of arthropod exoskeletons and fungal cell walls), *β*-glucosidase (G, break down cellulose, the structural component of plants), and phosphatase (P, breaks down organic monoesters for the mineralisation and acquisition of phosphorus). Cells were also counted at the end of the experiment and CO2 dissipation quantified as a single accumulative measure along the seven days of experiment. Full experimental details can be found in [32].

### Determination of classes

We computed all-against-all communities dissimilarities with Jensen-Shannon divergence [35], *D*_JSD_, and a transformation of the SparCC metric [36], *D*_SparCC_ (see Suppl. Material), and then clustered the samples following a similar approach to the one proposed in Ref. [18] to identify enterotypes. In the text, we call these clusters community classes. The method consists of a Partition Around Medoids (PAM) clustering for both metrics, with the function pam implemented in the R package cluster [63]. This clustering requires as input the number of output clusters desired *k*. We performed the clustering considering a wide range of *k* values and also computing the Calinski-Harabasz index (*CH*) that quantifies the quality of the classification, and selecting as optimal classification *k*_opt_ = arg max_*k*_(*CH*), shown in Fig. 2b. Processing of data and taxa summaries provided as Suppl. Material were generated with QIIME [64] and Phyloseq [65].

### Relation between community similarity and the sampling date and location

To investigate the relationship between the sampling location, the sampling date and the similarity in composition of bacterial communities, we performed analysis of the similarities of the communities grouping them with different criteria and testing if the similarities within groups were significantly different than the similarities between groups, using both *D*_JSD_ and *D*_SparCC_. We considered as grouping units one automatic spatial classification and two temporal classifications in which samples are joined in clusters depending on whether they were collected in the same day, or in the same month. Details for the spatial automatic classification and results for two other definitions of sampling sites (see Suppl. Material). We clustered the communities in spatial areas *A* of increasing sizes every order of magnitude, from 10 *m*^2^. to 100 *km*^2^, which we approximate considering spatial distances’ cut-offs of 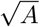 meters. We then computed the ANOSIM, MRPP and PERMANOVA tests (see Refs. [37, 38]) for each of the resultant classifications, using the R functions anosim, mrpp and adonis2, respectively (available in the R package vegan [66]) and assessing the significance with permutation tests (10^3^ permutations). To interpret the observed trends of these metrics we created synthetic distance matrices following different criteria, available in Suppl. Material.

### Structural equation modelling

Structural equation models [41] were built and analysed with lavaan (version 0.523) and visualized with semPlot R package [67, 68]. The modelling procedure was split into different stages detailed in Suppl. Material. First, a global model considering all data was investigated following several theoretical assumptions about the relationship between the functions, until a final model was achieved. Then, we looked for a second series of models in which it was possible to fit a different coefficient for each of the parameters in the global model, constraining the data into subsets corresponding to the community classes (i.e., six possible coefficients for each SEM pathway). Minor respecification of the model was performed (see Suppl. Material). We investigated whether altering the constraints on the models provided better fits, and penalized the models according to the number of degrees of freedom. The main criterion to accept a change was that the AIC of the modified model was smaller than the original model [69]. We verified that a several estimators were improved after any modification, including the RMSEA, the Comparative Fit Index and the Tucker-Lewis Index [70, 71].

Investigating causal relationships between endogenous and exogenous variables within the final specified model required controlling for confounding factors. For each pathway in the regression in the SEM model, we identified its adjustment set with dagitty [42]. We then performed a linear regression of each pathway adjusted by the confounding factors, adding a factor coding for the different classes. The coefficients obtained from the regression were estimated with respect to the reference class. Finally, we identified significant interaction terms between classes and the exogenous variable under investigation in the pathway. A significant interaction coefficient involving a given class was interpreted as a different performance of that class with respect to the reference class, and was therefore used to identify distinctive functional features of each class.

### Metagenomic analysis

Metagenomics predictions were performed using PiCRUST v1.1.2 [43] and quality controls computed (Suppl. Table. 8). A subset of genes appearing at intermediate frequencies was selected (Suppl. Fig. 17) and aggregated into KEGG pathways [72]. The mean proportion of genes assigned to a specific pathway was computed across communities belonging to the same class. Then we tested if the differences in mean proportions between classes were statistically significant using post-hoc tests with STAMP [73] (see Suppl. Material). To create Fig. 5 we visually inspected each post-hoc test and ranked the classes according with the number of pairwise tests in which they appeared significantly inflated (Suppl. Figs. 22–32). We qualitatively represent this ranking with circles of different sizes. Classes that do not appear inflated in any pairwise test in the pathway are not represented.

## Supporting information

Supplementary Methods and Results

## Acknowledgements

We acknowledge Damian Rivett for explanations about the experimental methods, and Matt Jones, Lara Durán-Trío and Yonathan Friedman for helpful discussions. We thank Andreas Steingötter from the ETH Seminar in Statistics for the support discussing the statistical methods. The research was funded by a European Research Council starting grant (311399-Redundancy) awarded to T.B. T.B. was also funded by a Royal Society University Research Fellowship. A.P.G. was also funded by the Simons Collaboration: Principles of Microbial Ecosystems (PriME), award number 542381.

