## Supplementary Methods and Results for "Community-level signatures of ecological succession in natural bacterial communities"

Alberto Pascual-García(1,2) and Thomas Bell(1)

November 29, 2019

(1) Department of Life Sciences. Silwood Park Campus. Imperial College London, Ascot, United Kingdom
(2) Current address: Institute of Integrative Biology. ETH-Zürich, Zürich, Switzerland

#### Contents

**1 Community composition and spatial autocorrelation** **2**
1.1 Sampling sites 2
1.2 Computation of $\beta-$diversity 2
1.3 Computation of ANOSIM, MRPP and PERMANOVA statistics 4
1.4 $\beta-$diversity classes 8
1.5 Examples of the spatial distribution of community classes. 13

**2 Structural equation modeling** **15**
2.1 SEM supplementary methods 15
2.2 SEM supplementary results 15

**3 Metagenomic analysis** **23**
3.1 Metagenomics prediction 23
3.2 PCA analysis 25
3.3 Post-hoc analysis of KEGGS's pathways 25

### 1 Community composition and spatial autocorrelation

#### 1.1 Sampling sites

In this work we analyzed 753 bacterial communities sampled from water-filled beech tree-holes in the South West of the United Kingdom [1]. These communities were sampled from different locations, with both the identifiers used by the field researchers and their popular names provided in Suppl. Tab. 1. Since the determination of sampling sites given by field researchers may be biased due to their subjective perception of the sampling areas, we performed an automatic clustering of the tree-holes looking for groups having significantly closer spatial distances within their own sampling area than with respect to other areas. We considered the GPS coordinates of each samples and calculated the distance between tree-holes through the haversine distance (R function DISTM, package GEOSPHERE). As we did in the determination of community classes (see Main Text), we used the PAM clustering for different $k$ values and then used the Rousseeuw-Silhouette index [2] (R function INDEX.S package CLUSTERSIM), obtaining clusters that closely reflect those determined by both the field-work researchers and the popular names attributed to the sites (see Suppl. Table 1). The median of the minimum, mean and maximum distance between tree-holes within the optimal sampling sites are 66.9 m., 539 m. and 1726 m., respectively.

#### 1.2 Computation of $\beta-$diversity

##### Dissimilarity metrics

We quantified the all-against-all communities dissimilarity metrics as follows. Calling $X_{ia}$ to the $N \times M$ matrix containing the number of 16S rRNA reads of OTU $i$ found in community (sample) $a$, we first computed the Jensen-Shannon divergence ($JSD$), which is a symmetrization of the Kullback-Leibler divergence:

$$JSD(a, a') = \sum_{i}^{S} \left( p_i \log \frac{2p_i}{p_i + q_i} + q_i \log \frac{2q_i}{p_i + q_i} \right),$$

where $p_i$ ($q_i$) is the relative abundance of OTU $i$ in the community $a$ ($a'$). We used $D_{\text{JSD}} = \sqrt{JSD}$ as a distance to compare the communities, that has metric properties thus fulfilling the triangle inequality, what makes it suitable for some clustering methods.

We next computed SparCC [3], which was originally developed to quantify correlations between OTUs $\rho_{ij}$ (see next section), and that we also applied to quantify correlations between communities, $\rho_{aa'}$, just by transposing the OTU table with the code provided in [3]. In a section below we verify that the SparCC is not affected by differences in the number of reads and that it has the interesting property of being unaffected by the OTUs marginal totals. For the clustering method discussed in the following, we transformed the correlation $\rho$ obtained with SparCC into a dissimilarity as $D_{\text{SparCC}} = \sqrt{2(1-\rho)}$.

##### Independence of $D_{\text{SparCC}}$ of the number of samples' reads

Here we verify that the SparCC score [3] is not affected by the differences in the number of reads across the samples, a common source of bias in the comparison of community composition metrics. To infer $\rho_{aa'}$ we consider the vectors $\bar{y}_a = \bar{X}_a./\bar{n}$, where $\bar{X}_a = \{X_{1a}, ..., X_{Na}\}$, $\bar{n} = \{n_1, ..., n_N\}$ are the OTUs totals (i.e., $n_i = \sum_a X_{ia}$), and ./ the element-wise division. Next, a matrix containing the dependencies between the two communities $a$ and $a'$ is computed:

$$t_{aa'} = \text{var}\left(\log\left(\frac{\bar{y}_a}{\bar{y}_{a'}}\right)\right) = \text{var}\left(\log\left(\frac{\bar{X}_a}{\bar{X}_{a'}}\right)\right),$$

where the cell quantities $t_{aa'}$ are independent of the OTUs marginal totals. From this matrix, SparCC develops an heuristic to approximate the desired correlations $\rho_{aa'}$. We may still think that this approximation may generate spurious dependencies arising from the different number of reads of the samples.

To address this question let us consider a sample $\bar{X}_a$ with $m_a$ observations taken from an arbitrary probability distribution, and then another sample of the same distribution $\bar{X}_{a'}$ with $m_{a'}$ observations, such that $m_a \neq m_{a'}$. We want to verify that, when we compare these two samples against another sample $\bar{X}_b$ generated from a different probability distribution, the quantities $t_{ab}$ and $t_{a'b}$ are the same. Since the samples $\bar{X}_a$ and $\bar{X}_{a'}$ are generated from the same distribution it holds that $\bar{X}_{a'}/m_{a'} = \bar{X}_a/m_a$, and hence $\bar{X}_a = c\bar{X}_{a'}$, with $c = m_a/m_{a'}$. It follows that

| Autom. clustering Id. | Num. of samples | Researchers | Popular name |
|---|---|---|---|
| **1** | **147** | **AE** | **Ashridge Estate** |
| **2** | **72** | **BB** | **Burham Beeches** |
| **3** | **8** | **BCW** | **Birchcleave Wood** |
| **4** | **18** | **BCW** | **Ash Copse** |
| **5** | **24** | **BW** | **Bisham Woods** |
| **6** | **3** | **BWd** | **Beech Wood** |
| **7** | **16** | **HW** | **Hunts Woods** |
| **8** | **43** | **LW, SNG, SP, WGP, WiW** | **Long Wood, Sunningdale, Silwood Park, Windsor Great Park1, Greenbroom Covert1** |
| **9** | **77** | **OWW, WF, WGP, WiW** | **Old Windsor Wood, Windsor Forest, Windsor Great Park2, Greenbroom Covert2** |
| **10** | **23** | **PH** | **Pullingshill Wood** |
| **11** | **15** | **SAC** | **StAlbans City** |
| **12** | **25** | **WEL** | **Combe Hill** |
| **13** | **15** | **WP** | **Waterers Park** |
| **14** | **186** | **WYC, WYD, WYM, WYT** | **Wytham Woods** |

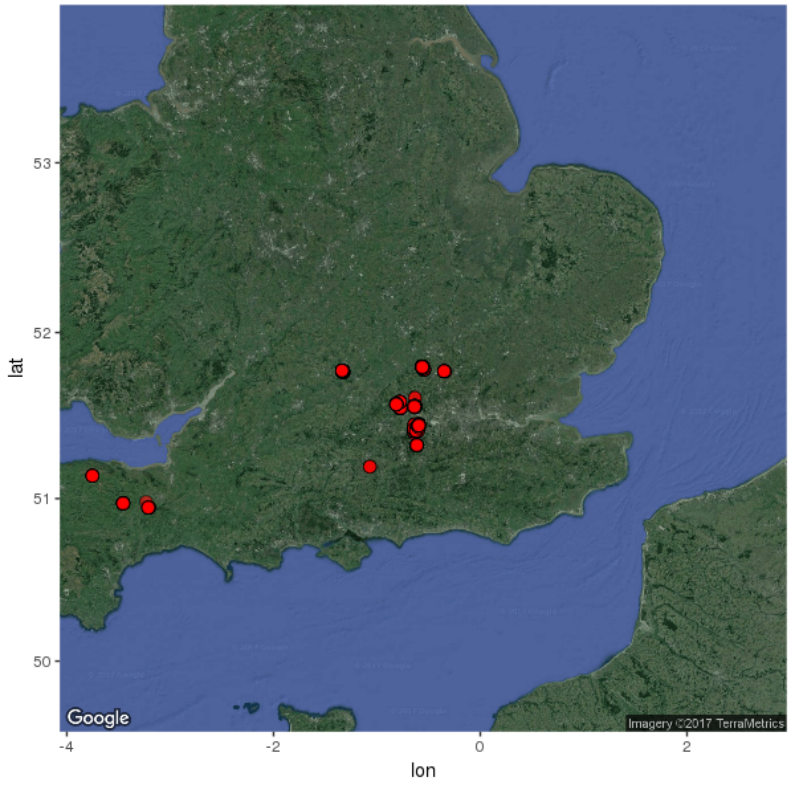


Table 1: **Sampling sites.** (Above) Relation between the three definitions of sites considered in this work. Labeling of the sites provided by field researchers, may depend on biased perceptions of the space or the specific day in which the sample were collected, and thus an automatic clustering using spatial distances was made (first column, see Methods in the main text). It is mostly consistent with the researchers labelling, except for the areas around Windsor Great Park, that were more heavily sampled and the automatic clustering join them together, while researchers considered them different locations. Another example are locations like Birchcleave Woods, that should be likely splitted but the automatic clustering joins them, possibly affected by the fact that are very far away from all other locations. (Below) Overview of the sampling locations. Most of the samples were collected in sorrounding areas of Silwood Park campus and Oxfordshire (centre of the map), with a second distant cluster in the South West of UK.

$$t_{ab} = \text{var}\left(\log\left(\frac{\bar{X}_a}{\bar{X}_b}\right)\right) = \text{var}\left(\log\left(\frac{c\bar{X}_{a'}}{\bar{X}_b}\right)\right) = \text{var}\left(\log(\bar{X}_{a'}) - \log(\bar{X}_b) + \log(c)\right) =$$
$$= \text{var}\left(\log(\bar{X}_{a'}) - \log(\bar{X}_b)\right) = \text{var}\left(\log\left(\frac{\bar{X}_{a'}}{\bar{X}_b}\right)\right) = t_{a'b}.$$

#### 1.3 Computation of ANOSIM, MRPP and PERMANOVA statistics

##### Supplementary results for observed data

We classified samples at different distance thresholds using the haversine distance matrix, and clustering the samples with single linkage clustering implemented in the function HCLUST in the package STATS. We then stopped the clustering at every order of magnitude of the cluster's area $A$, from 10 to $10^9$ $m^2$., which can be approximated stopping the clustering when the threshold in which we join two clusters is $\sqrt{A}$. This leads to 10 classifications of samples in spatial clusters at different scales. We then performed three types of statistical tests: ANOSIM (Analysis of Similarities), MRPP (Multi Response Permutation Procedure) and PERMANOVA (Permutational Multivariate Analysis of Variance Using Distance Matrices) to evaluate if the $\beta-$diversities of the samples within clusters of a given classification were significantly different from those belonging to other clusters. Significance was estimated with $10^{-3}$ permutation tests. In Suppl. Fig. 1 we show the significance of ANOSIM for the spatial clustering at the shortest and largest distances, where the values of the statistics differ in one order of magnitude. Similar results were found for the other metrics at all classifications.

In addition, we provide results for a PERMANOVA test in Suppl. Fig. 2. In contrast with the results found for ANOSIM and MRPP, this statistic do not depict any clear trend across the different scales. To interpret the discrepancy between the different metrics, in the next section we study the behaviour of these metrics with synthetic data.

##### Analysis of synthetic data

To understand in more detail the trends found for the different statistical measures, we designed an analysis in which we tested the performance of these quantities with synthetic data. We created synthetic distances matrices of 1000 elements that are regularly clustered in the same number of levels than the observed data. The distances within and between clusters are determined with different criteria.

The first criteria considers distances that increase constantly across increasing scales (i.e. larger clusters). This is motivated by the observation of the MRPP trend in real data: the $\delta$ estimator computes the mean distance within the clusters and we observe a constant increase across scales. To implement this test we homogeneously divided the interval (-1,1) (which represents the possible values of the correlations provided by SparCC) by the number of levels considered, and we transformed it into a distance (see Section 1.2). Each value will represent the mean of a Gaussian distribution with a given standard deviation, around which random values will be generated for each level. We then created different matrices with increased standard deviation, from 0.1 to 1. An increase in the standard deviation can be interpreted in terms of increased taxa dispersion, because the probability that two distant locations are more similar than communities closer in space will be higher. An illustration of matrices obtained following this procedure is shown in Suppl. Fig. 3.

A good estimator should be able to reject the null hypothesis stating that there are no differences between the distances within– and between–clusters, as these differences exist by construction. In addition, from the point of the additional information it returns, if the standard deviation is small, it should be roughly constant for any of the tests across scales because, even if the mean distances decrease, the ratio between the mean within- and between-cluster distances are kept constant. Nevertheless, when the dispersion increases, the distributions generating a larger number of distances (which are those found for larger clusters, emulating larger distances scales) will also generate a larger number of distances lying at other levels. This should generate some decrease in the value of the estimator.

We analyzed the performance of the three statistics for these matrices for different values of the standard deviation. All three metrics are able to identify in all cases that there are significant differences across levels ($10^3$ permutation tests), and they are therefore capable to reject the null hypothesis that there are no differences between the distances within and between clusters. We anticipated that the MRPP estimator ($\delta$) computes the overall weighted mean of within-group means of the pairwise distances among clusters, and hence it must depict a perfect linear trend with respect the mean distances generated independently of the standard deviation (see Suppl. Fig. 4, left column). Therefore, $\delta$ help us to control the structure built on the distances matrix, but it

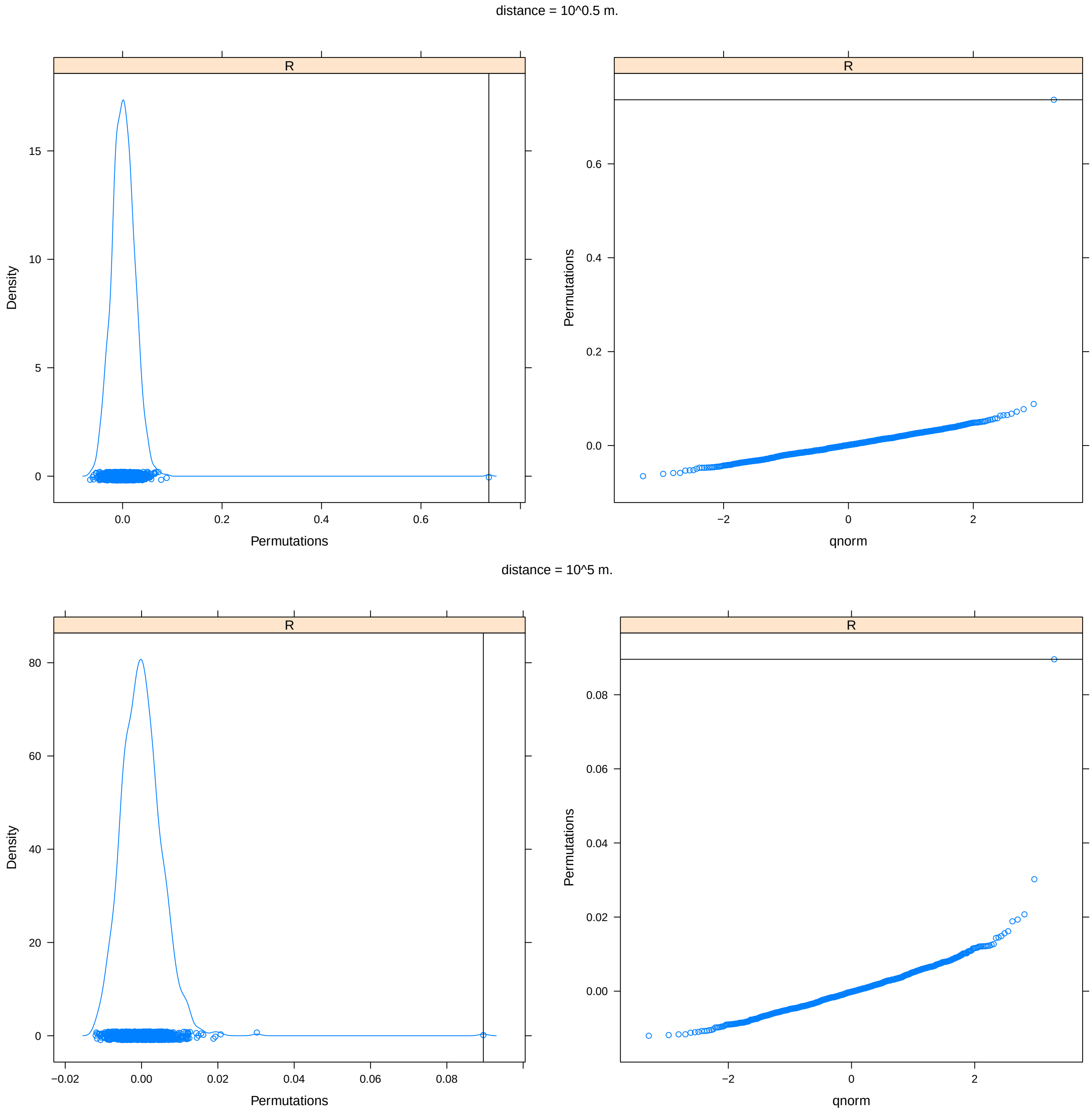


Figure 1: **ANOSIM permutation tests.** Density of the ANOSIM values for permuted $\beta-$diversity matrices (left boxes) and its quantile normalized values (right boxes) for communities clustered at spatial distances thresholds $\sqrt{10}$m. (top) and 100km. (bottom).

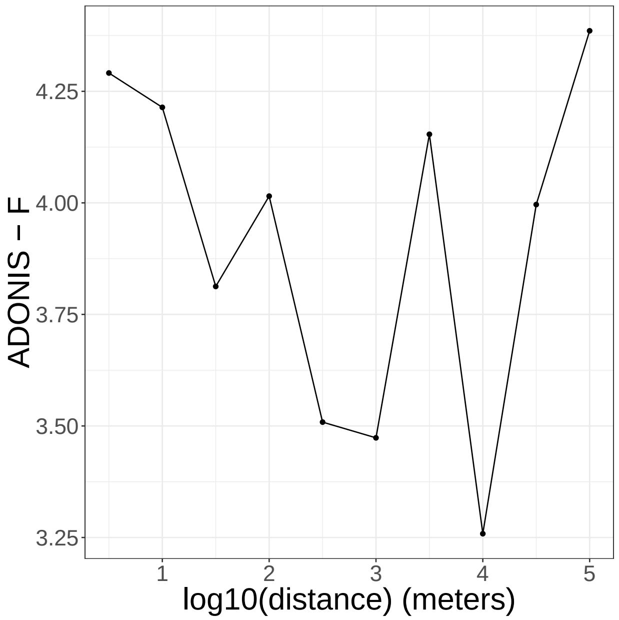


Figure 2: **PERMANOVA analysis**. Values of the $F$ statistics for the classifications found at the different spatial distances. The intermittent behaviour contrast with the continuous changes found for ANOSIM and MRPP shown in Main Text.

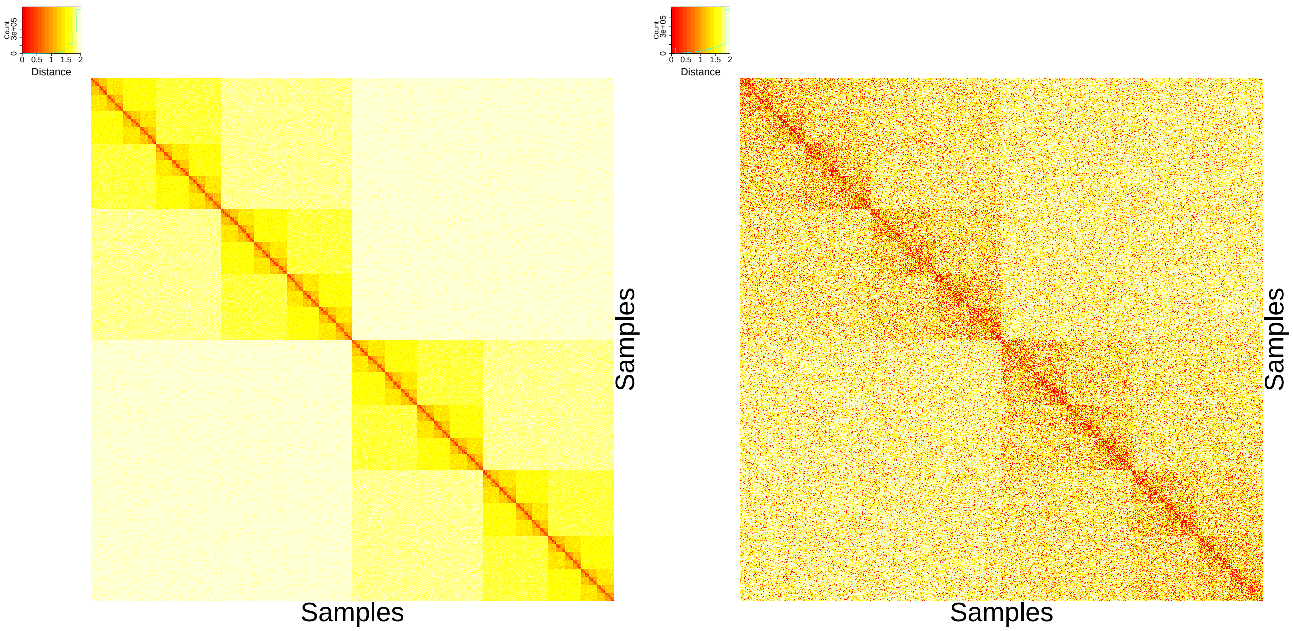


Figure 3: **Synthetic distances matrix** generated with an increasing mean for larger clusters with a standard deviation equals to 0.1 (left) and to 1 (right).

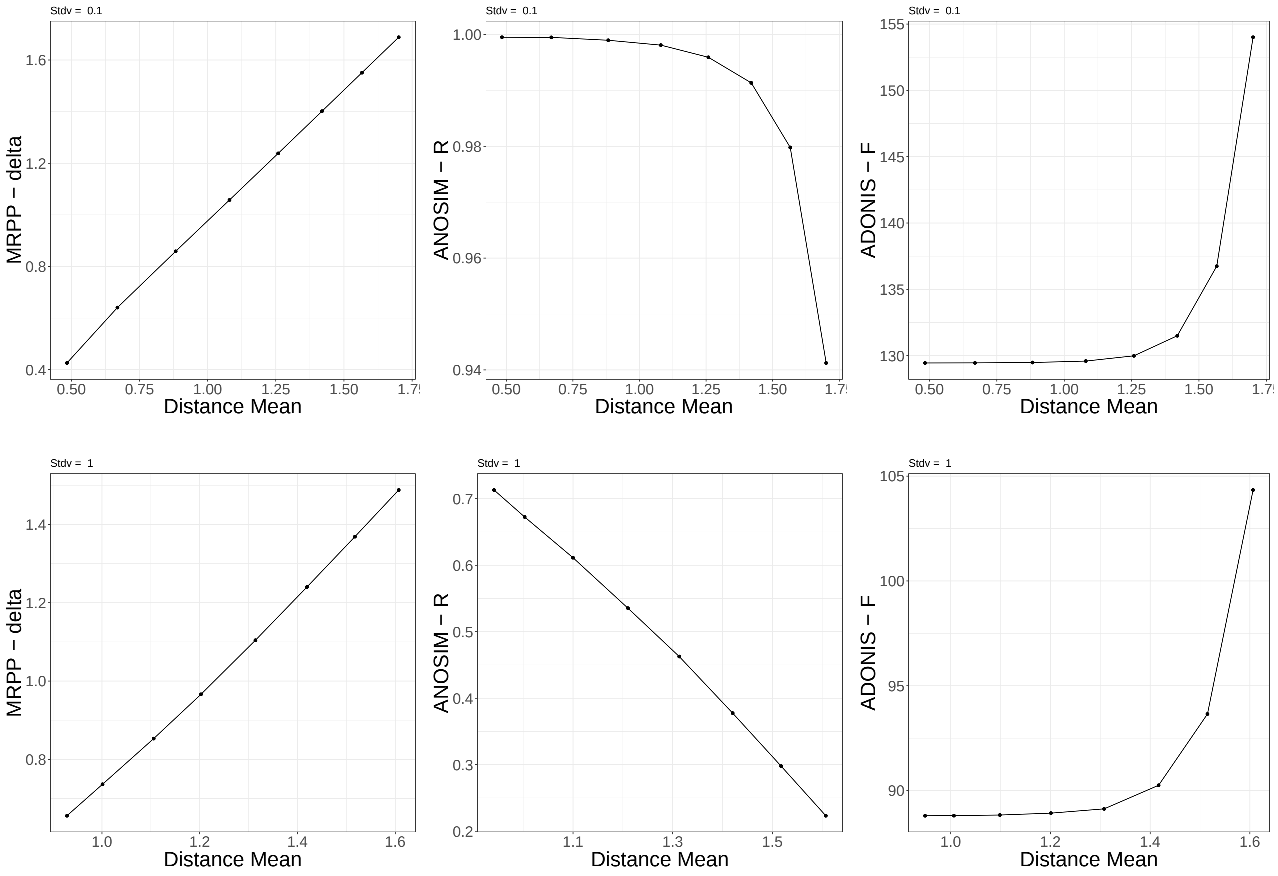


Figure 4: **Statistics of synthetic matrices** with distances between blocks increasing linearly across levels. (Left column) $\delta-$value of the MRPP test (middle) $R$ statistics of the ANOSIM test and (right) $F$ statistics of a PERMANOVA test for $\sigma = 0.1$ (top row) and $\sigma = 1$ (bottom row), against the mean distance within each level.

is insensitive to any increase in the dispersion. For ANOSIM, we observe a decrease in the estimator ($R$) with increasing distances. This decrease is small in absolute value (from 1 to 0.94, note that $R \in [-1, 1]$) for small standard deviations ($\sigma = 0.1$, see Suppl. Fig. 4 middle-top), and it increases for increasing $\sigma$, resulting very steep for $\sigma = 1$. Therefore, the performance of this statistics follows the expected behaviour for a "good" estimator outlined above for the problem we aim to investigate. For the $F$ statistics of the PERMANOVA test, it increases with respect to the man distances generated for any value of $\sigma$, and it just changes the order of magnitude of the estimator, which is in all cases very high (see Suppl. Fig. 4, right column).

Since the PERMANOVA statistics for the observed data shows an intermittent behaviour (see Suppl. Fig. 2), we investigated this pattern generating matrices following a second criteria. We generated matrices for which the mean distance increases, with additional abrupt decreases of the distances for some levels, for which we generated distances with the parameters of the distribution of the first (lowest distances) level (see Suppl. Fig. 6). Again, we confirmed that the $\delta$ value of MRPP essentially quantifies the mean distance of the clusters (see Suppl. Fig. 6 left), with both ANOSIM and PERMANOVA estimators being able to retrieve the abrupt introduction of a decrease in the distances, and the ANOSIM estimator being the only one detecting an increase in the standard deviation (comparison of top and bottom rows in Suppl. Fig. 6). The PERMANOVA estimator seems to be insensitive to these changes and the fact that it again increases for the last levels, which is were the clusters are larger and hence have a much higher number of distances, inclines us to think that the intermittent behaviour observed in real data comes from the heterogeneity in sizes of the clusters, since in the synthetic data all clusters are homogeneous.

Interestingly, it has been previously claimed in the literature that the PERMANOVA statistics is a more robust estimator than the ANOSIM method [4], being the latter more sensitive to environmental changes [5]. For the problem we are handling, it seems that this sensitivity is a desirable property to disentangle changes in

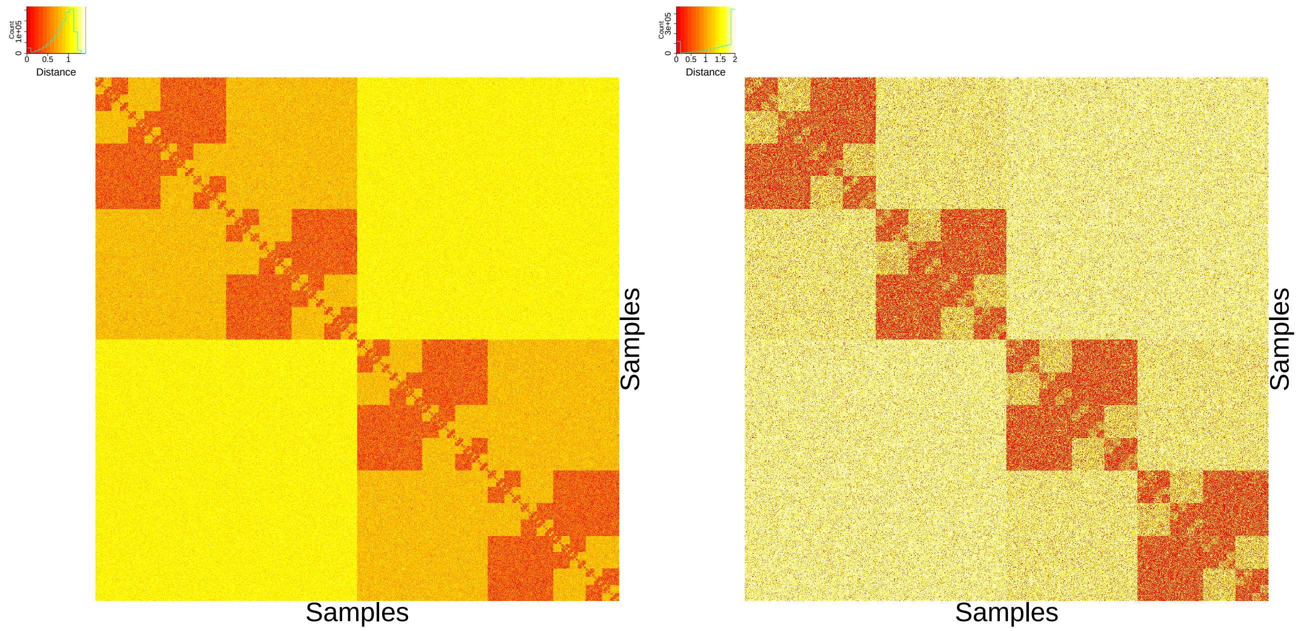


Figure 5: **Synthetic distances matrix** generated with an increasing mean for larger clusters and an intermittent decrease of the distance every two levels to the lowest distance values, with a standard deviation equals to 0.1 (left) and to 1 (right).

mean and variance of the distances across the different scales, with the analysis we conducted pointing towards and increase in the mean of the $\beta-$diversity distances with respect to an increase in the spatial distance of the samples, and a high dispersion (reflected in the high variance needed to obtain the strong distance-decay trend observed in real data).

To get a quantitative understanding of the implications of the observed ANOSIM decay, we conducted an analysis for the matrices built with the first criteria (constant increase of the mean distances for larger clusters, Suppl. Fig. 4), and we estimated which is the probability $P$ that a distance generated in a classification at a given level $\alpha$ is lower or equal than the distance generated in the first (baseline) level (labeled 0: the one with the lowest distances, i.e. $P(d_{ij}^{\alpha} \leq d_{ij}^{0})$ with $d$ being the $\beta-$diversity distance). This quantity estimates how likely it is that two communities distant in space are as similar as those very close in space. While for low levels of dispersion this probability is very small for levels far from the baseline level (Suppl. Fig. 7 left), to get a synthetic trend comparable with the observed data (i.e. the range of maximum and minimum values of the curve are similar), we need to generate distances with large standard deviations (see e.g. $\sigma = 1$ Suppl. Fig. 4 middle column, second row). In this situation, $P(d_{ij}^{\alpha} \leq d_{ij}^{0})$ is high even for the most distant levels, with values between 0.03 and 0.10 for the last three levels. For the observed data, this would imply that samples separated by several kilometers (more than one hundred for the last level) have a noticeable probability of being more similar than those located few meters away. We cannot foresee any transport process justifying this observation, what led us to investigate the existence of similar environmental conditions in distant locations as an alternative hypothesis. Scripts for the analysis performed in this section are available at https://github.com/apascualgarcia/TreeHoles_descriptive.

#### 1.4 $\beta-$diversity classes

##### Description and determination of $\beta-$diversity classes

Using the $\beta-$diversity metrics $D_{\mathrm{JSD}}$ and $D_{\mathrm{SparCC}}$, and following the automatic classification pipeline explained in the Main Text, we found six community classes. In Suppl. Fig. 8 we show the results of the Calisnki-Harabasz index versus the number of final clusters in the classification for $D_{\mathrm{JSD}}$.

In addition, in Suppl. Fig. 9 we show the relative abundance of the most important OTUs for the classes obtained with $D_{\mathrm{SparCC}}$, which are those used in most of the analysis presented.

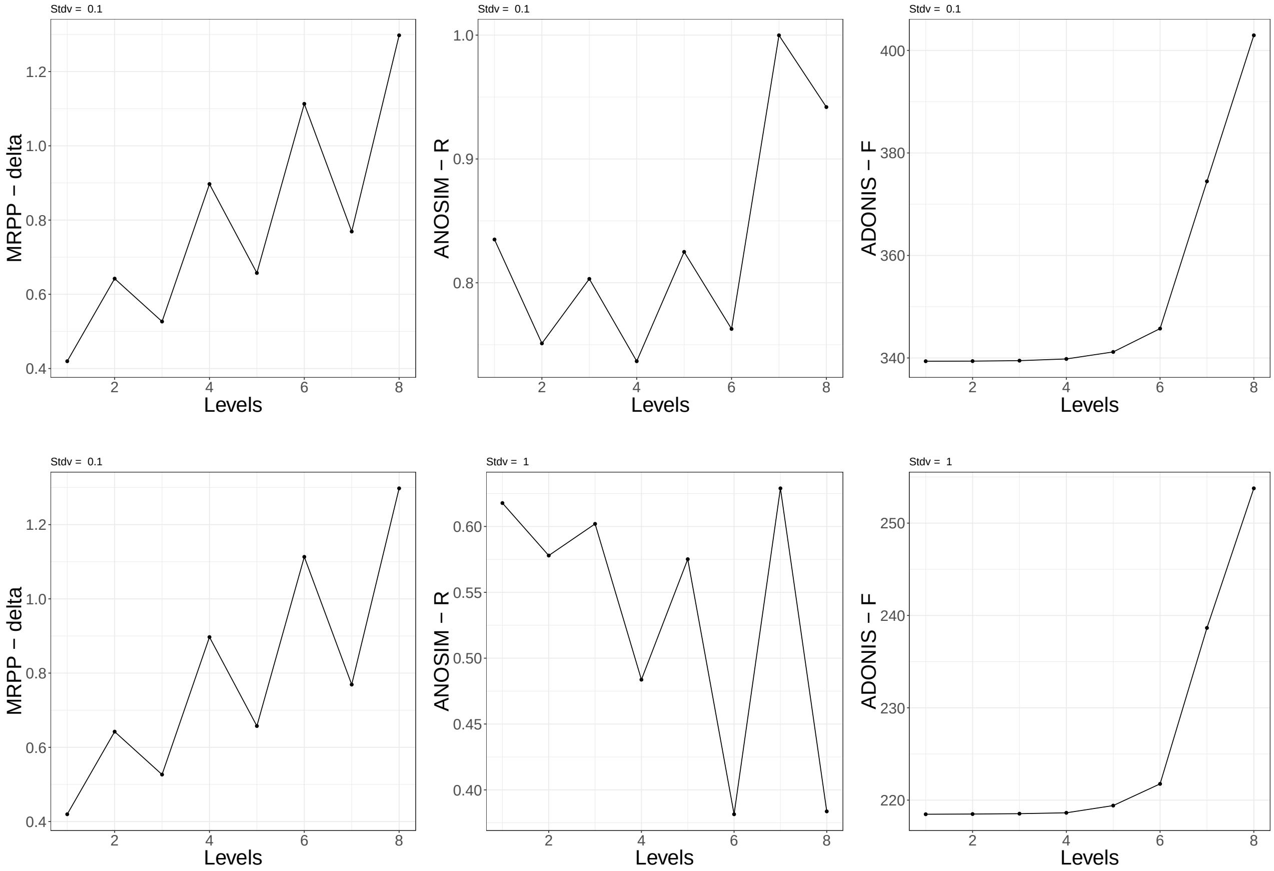


Figure 6: **Statistics of synthetic matrices** with distances between blocks increasing linearly across levels and with intermittent drops in the distance. (Left column) $\delta-$value of the MRPP test (middle) $R$ statistics of the ANOSIM test and (right) $F$ statistics of a PERMANOVA test for $\sigma = 0.1$ (top row) and $\sigma = 1$ (bottom row), against the rank of each level.

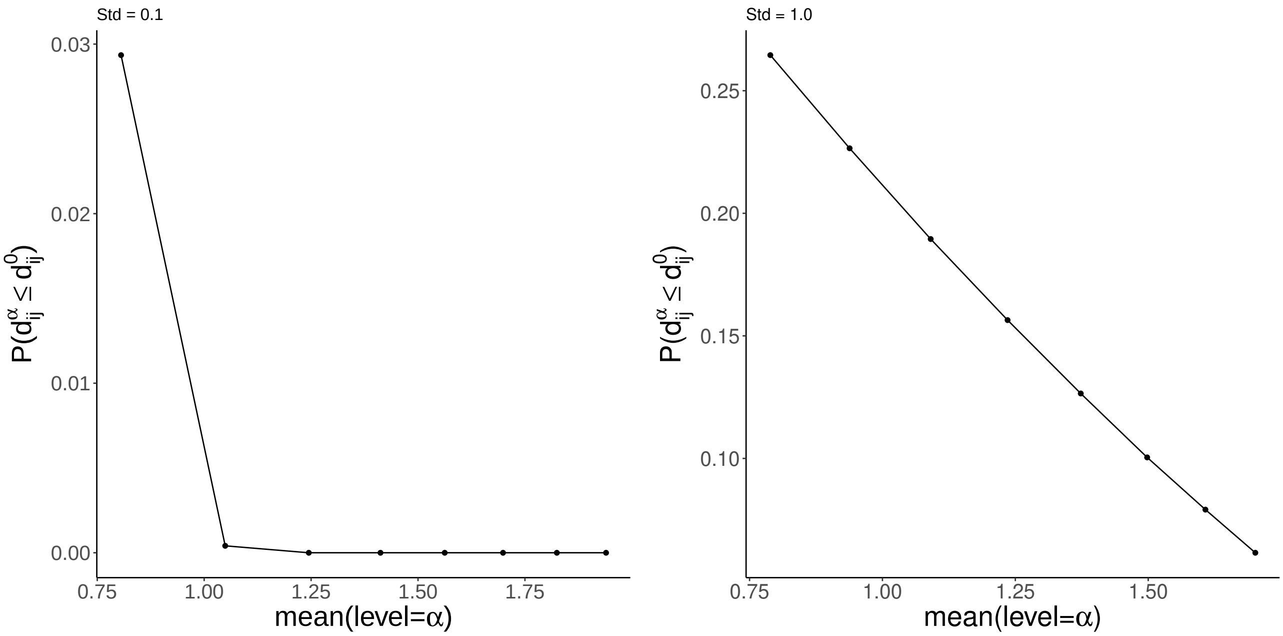


Figure 7: **Overlap between the distances generated at the different levels $\alpha$ and the level with the lowest distances** (labeled 0), against the mean of the distances of each level, for $\sigma = 0.1$ (left) and $\sigma = 1$ (right).

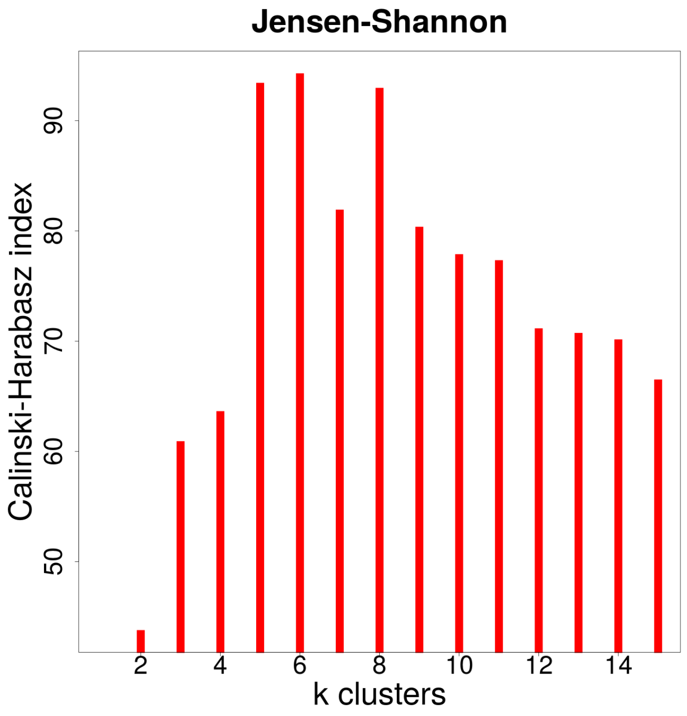


Figure 8: **Number of classes obtained with $D_{\text{JSD}}$.** Calinski-Harabasz index measured for the PAM agglomerative clustering with different number of final clusters.

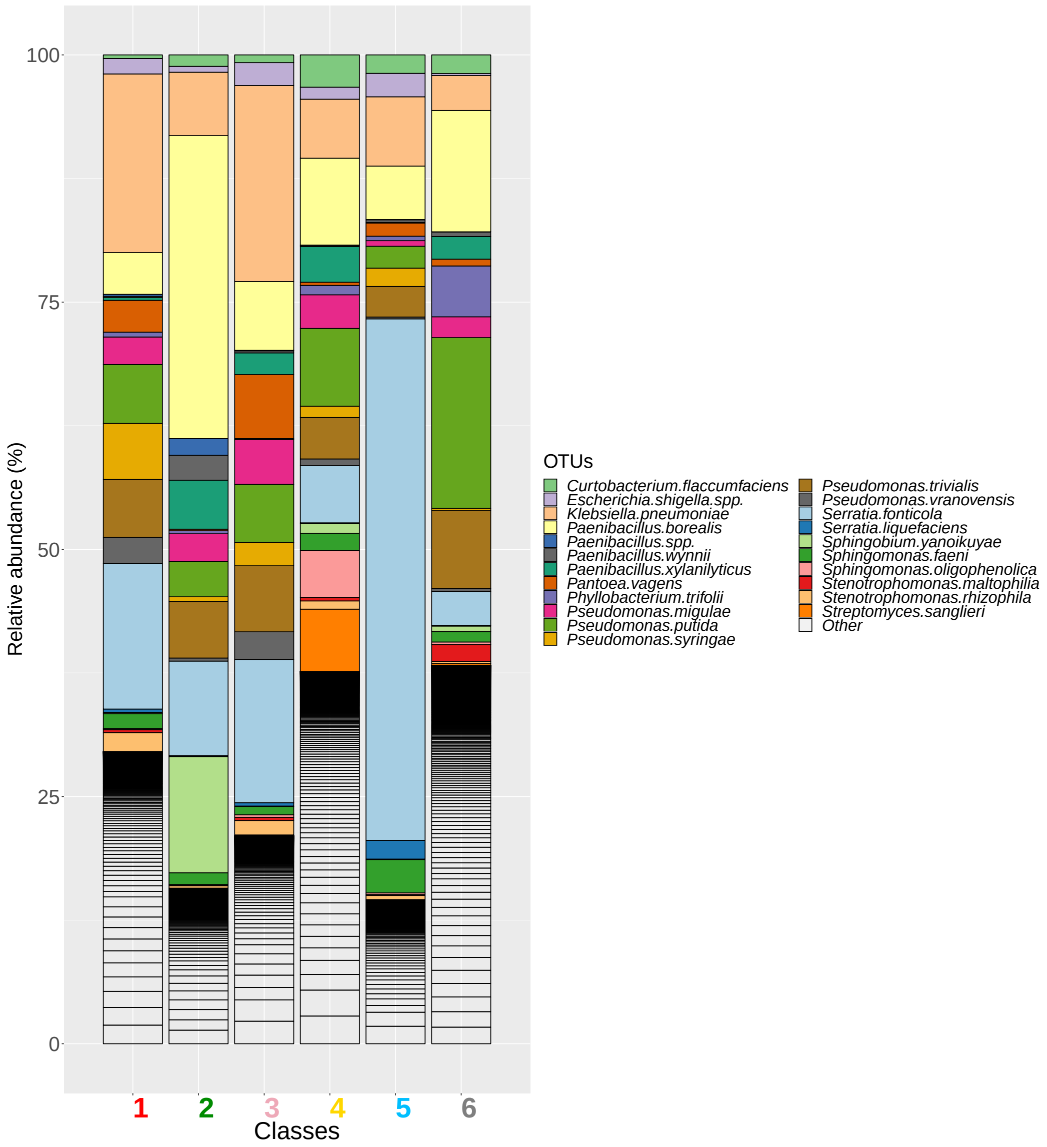


Figure 9: **OTUs composition across the six classes.** Taxa among the 15 highest abundant in any of the classes are highlighted with the remainder classified as "Other".

| Class | Num. of communities | OTUs Num. | Mean num. of reads | Effective Biodiversity |
|---|---|---|---|---|
| 1. Red (Klebsiella) | 194 | 151 (21) | 16785 (530) | 13.7 (5.5) |
| 2. Green (Paenibacillus) | 99 | 127 (25) | 17289 (905) | 4.4 (2.4) |
| 3. Pink (Klebsiella) | 124 | 128 (16) | 16891 (651) | 9.5 (2.9) |
| 4. Yellow (Pseudomonas) | 77 | 127 (26) | 16556 (1677) | 6.2 (4.6) |
| 5. Blue (Serratia) | 100 | 125 (22) | 17276 (1339) | 5.1 (2.6) |
| 6. Grey (Pseudomonas) | 86 | 143 (33) | 18606 (4914) | 17.1 (12.0) |

Table 2: **Properties of the different classes.** The table summarizes the number of communities, and the mean number of OTUs found at 97% 16S rRNA similarity, after excluding rare species and samples with low number of reads (see Methods in Main Text), the mean number of reads as a proxy of the species abundances and the mean effective biodiversity. Values within parenthesis stand for standard deviations.

##### Properties of the $\beta-$diversity classes

Properties of the $\beta-$diversity classes are summarized in Suppl. Table 2. The mean number of reads are comparable across classes. The effective diversity, computed as $EB = \exp(S)$, with $S = -\sum_i f_i \ln f_i$, where $f_i$ is the relative abundance of species $i$ in the sample, are low, reflecting the dominance of few species. Note that we excluded species with less than 100 reads across the 680 samples, approximately representing half of the OTUs. This facilitates the identification of well separated classes, but it is a criteria that should be analysed for each dataset observing the influence in the number of clusters of removing species.

##### Comparison between the $\beta$-diversity classes the sampling date and the sampling location.

We investigated if the classifications obtained with $D_{\mathrm{JSD}}$ and $D_{\mathrm{SparCC}}$ were significantly similar, and their relation with classifications of samples according to their sampling date or their sampling location. We used five indexes (Rand, Fowlkes and Mallows, Wallace 10, Wallace 01 and Jaccard) to compare the classifications obtained, as implemented in the R PCI function of the PROFDPM package [6]. Unfortunately, it is difficult to determine if a given index has a significant value, and thus we developed the following strategy. Firstly, we obtained for each index $x$ the mean $\bar{x}_B$ and standard deviation $\sigma_B$ of the index when the samples are bootstrapped $10^3$ times. This will give us a confidence interval around the observed classification. To continue with, we shuffled the identifiers relating each sample with one of the classifications, what would randomize the mapping between samples and the classification while maintaining the relative number of samples for each class. We perform the shuffling $10^3$ times, calculating the index for each shuffling, and retrieving the maximum value obtained $x_R$, what will give us the maximum value that can be obtained randomly. We aim to verify that the random value is significantly different than the bootstrapped distribution. Hence, given the observed value $\hat{x}$, we computed two z-scores:

$$\hat{z} = \frac{\mathrm{abs}(\hat{x} - \bar{x}_B)}{\sigma_B}; \; z_R = \frac{\mathrm{abs}(x_R - \bar{x}_B)}{\sigma_B},$$

where $\hat{z}$ reflects the bias of the observed value, and $z_R$ the deviation of the random value with respect to the bootstrapped distribution. All values must be negative, and we took the absolute value for simplicity. We verified that there is no significant bias, being all the $\hat{z}$ values close to zero. We considered that the observed value $\hat{x}$ is significant if $z_R > 2$.

We compared these four classifications in an all-against-all basis confirming that the classifications found with $D_{\mathrm{SparCC}}$ and $D_{\mathrm{JSD}}$ are significantly similar and that they do match better with respect to the day of collection than with the sampling areas (see Suppl. Table 3 and Suppl. Methods). $D_{\mathrm{SparCC}}$ seems to capture again better the similarities than $D_{\mathrm{JSD}}$, being its values with respect to space and time classifications remarkably higher, and hence it is the one selected to present most of the analysis. It was finally noted that the classification of classes given by clustering of the $D_{\mathrm{SparCC}}$ and $D_{\mathrm{JSD}}$ metrics and selection of an optimal threshold, led to much higher ANOSIM values, which suggests that these classes captured local conditions occurring even at distant locations (see Fig. 1 in Main Text).

In addition, in the analysis of the similarity of the communities shown in Fig. 1 in Main Text, we included a value labeled as "Sites" that describes the ANOSIM estimation for the sampling sites obtained with the above

| Comparison | Rand | Fowlkes-Mallows | Wallace 10 | Wallace 01 | Jaccard |
|---|---|---|---|---|---|
| $D_{\text{SparCC}}$ Vs. Day | 2.42 | 9.02 | 6.80 | 7.77 | 8.51 |
| $D_{\text{SparCC}}$Vs. Space | 2.31 | 5.58 | 4.60 | 4.77 | 5.40 |
| $D_{\text{JSD}}$ Vs. Day | 4.62 | 4.79 | 3.65 | 4.52 | 4.49 |
| $D_{\text{JSD}}$ Vs. Space | 4.39 | 4.53 | 3.41 | 4.66 | 4.26 |
| $D_{\text{JSD}}$ Vs. $D_{\text{SparCC}}$ | 9.13 | 10.37 | 9.69 | 9.12 | 9.29 |

Table 3: **Comparison of classifications.** Z-score of different indexes (columns) to compare classifications (rows) when one of the classifications is randomized, and the bootstrapped distribution of the correspondent index is considered (see Suppl. Methods). Higher the Z-score more similar the classifications. We considered significant values higher than 2, being the classifications obtained with $D_{\text{JSD}}$ and $D_{\text{SparCC}}$ the most significantly similar, and being both more similar to the classification related with date of sampling (Day) than with the site as defined by the automatic classification (Space).

| | Day | Month | Automatic | Popular | Researcher | Classes |
|---|---|---|---|---|---|---|
| $D_{\text{SparCC}}$ | 0.42 | 0.34 | 0.26 | 0.27 | 0.35 | 0.60 |
| $D_{\text{JSD}}$ | 0.28 | 0.34 | 0.23 | 0.23 | 0.24 | 0.59 |

Table 4: **ANOSIM $r$ statistics** for two temporal classifications (samples collected at the same day or at the same month), three definitions of sampling sites ("Automatic", "Popular" and "Researcher", see Methods), and the intrinsic classes obtained with automatic clustering for both $D_{\text{SparCC}}$ and $D_{\text{JSD}}$.

automatic procedure. In Table 1 we additionally show the results for the other two sites schemes, together with the results found if the communities are clustered according to the day or the month of sampling collection.

#### 1.5 Examples of the spatial distribution of community classes.

In Suppl. Fig. 10 we show several examples of the distribution of the classes in the space. While some communities belong to the same classes within the same site (upper rows) other sites host several classes (lower rows).

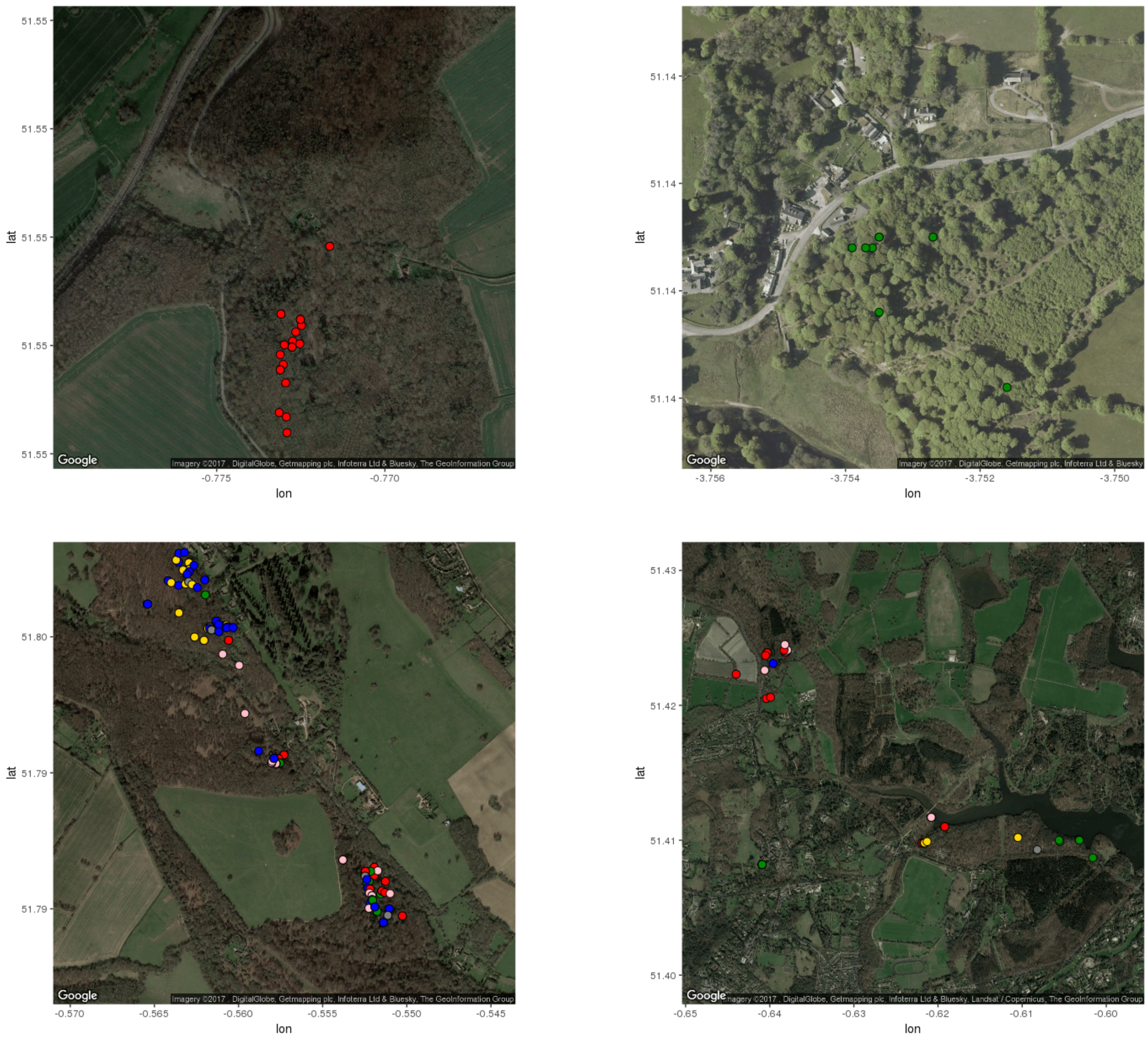


Figure 10: **Examples of spatial distribution of the classes.** (Above) Examples of sites were only one specific class is observed, Bisham Woods (left) and Birclave Woods (right). (Below) Larger scale examples were it is possible to observe different sites: Ashridge Estate (left) and Windsor Great Park (right) Class 1 = red (reference); Class 2 = green (Paenibacillus); Class 3 = pink (Klebsiella); Class 4 = yellow (Streptomyces); Class 5 = blue (Serratia); Class 6 = grey (P. putida).

### 2 Structural equation modeling

#### 2.1 SEM supplementary methods

More than 200 models were analyzed and visualized with LAVAAN (version 0.523) and SEMPLOT R packages [7, 8]. The results of the fits are available in https://github.com/apascualgarcia/TreeHoles_descriptive. The models were conducted by averaging the replicas, given the high intrinsic noise of these experiments. We divided the problem into different stages. In the first exploratory stage, we investigated the basic relationships between the variables and whether it would be convenient to group them into latent variables, leading to a first satisfactory model. In the second stage, we introduced the definition of the different classes which, given the large number of new parameters introduced, led to a worse model, as determined by the AIC value. Therefore, we respecified the model, finding two relevant modifications: the deletion of a pathway between P and Cells and the inclusion of all the correlations between exoenzymatic variables. The third stage investigates constraints in the parameters for the different variables across classes.

We used different criteria to determine which is the best model in this progression of models. In all cases, we considered the $R^2$ values of the exogenous variables, and the significance of the coefficients inferred. Then, a battery of indexes [9, 10, 11] such as the RMSEA together with a 90% CI, the Tucker-Lewis index, the Comparative Fit Index, and the AIC and BIC indexes. We observed that all the indexes change consistently. We also observed that the CI of the RMSEA is the most sensible indicator when the fits are close to the final model selected.

To investigate the relevance of community classes in the model, we started estimating one parameter per path and class leading to a "free" (of constraints) model, and then we investigated how introducing constraints (e.g. forcing the coefficients of a given pathway to be the same for some combination of classes) led to a better model, following the criteria explained above. We should note that we have $\sum_i \binom{6}{i} = 2^6 - 1 = 63$ ways to constrain each coefficient (e.g. class 1 and 2 have one coefficient and 3 to 6 another coefficient) and the order in which the coefficients are constrained may affect the results, therefore there is a combinatorial explosion in the number of models to be explored what makes an exhaustive analysis unfeasible. In addition, automatic searches in modeling specification are not recommended [12], and therefore we decided to develop the following heuristic. The rational behind the heuristic is that it will be more likely to find a better model if we constrain first those coefficients that are more similar, than if we constrain those that are very dissimilar. We started considering the model in which the parameters for each coefficient are estimated independently for each community class (the "free" model) and we proceeded as follows:

- For each coefficient, we computed a distance matrix between the standardized coefficients obtained for each class using an euclidean distance, and we clustered them using average linkage, illustrated in Fig. 11 (STATS R package).
- We tested the free model with respect to a model in which all the classes have the same value. The latter model corresponds with the last step of the clustering. If the free model was not rejected, we accepted the estimation of one coefficient for each of the classes.
- If the free model was rejected, we contrasted the model with a single coefficient with a model having two coefficients, constraining the classes according to the clusters found in the next-to-last step of the clustering (in Fig. 11 one coefficient will be assigned to classes 4 and 5 and another to classes 1, 2, 3 and 6). We repeated the test and, if the model with one constraint was rejected, we iterated the analysis considering a model with three coefficients (in Fig. 11, we would estimate the coefficient of class 6 independently). We finished the iteration when no further improvement was achieved.

We performed tests in which we randomly constrained the parameters in different groups, which never led to better results than the best model found following this scheme, so we believe that the procedure leads to close-to-optimal models.

#### 2.2 SEM supplementary results

##### Description of the functional trends

We investigated the functional capacities of these communities in laboratory environments (see Methods in Main Text), relative to their splitting into classes. While there is a large body of literature relating microbial functions

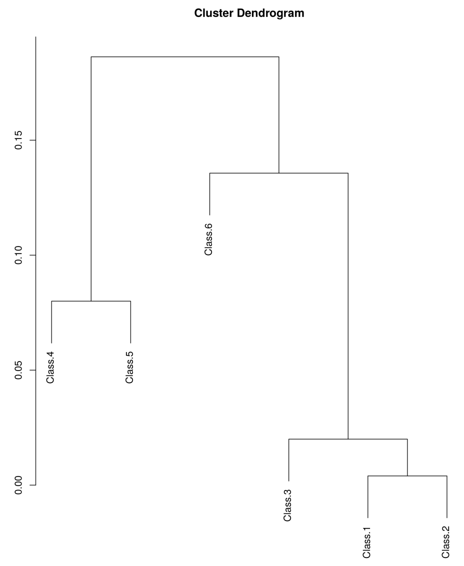


Figure 11: **Example of clustering of coefficients** estimated independently for each community class in a SEM for the partial regression Cells $\sim$ ATP.

such as exoenzymatic activities or respiration to microbial productivity [13], the differences in the bacterial composition are rarely explicitly interrogated, but rather assumed by the specific location in which the samples were taken. This makes difficult to individuate if the patterns found are due to actual differences in community composition, or to the different activity that these enzymes have under different environmental conditions. In this study, the environmental conditions in which bacterial communities are grown are the same, and therefore differences in functioning should be attributed to communities composition.

In Suppl. Fig. 12 we show linear regressions between the log-transformed variables, finding marked differences in some of the variables for the different classes. For instance, class 5 (blue) had a not-significant relation between cell count and CO2 produced, that may reflect a strictly anaerobic performance. As another example, class 6 (grey) had a large number of points with low cell count and large CO2 production, consistent with the hypothesis we propose in the Main Text, stating that these communities are poorly adapted to this environment. Indeed, given that the number of cells is lower than the starting number introduced in the experiment ($10^4$) this signal might be due to the release of endoenzymes after death, which is known may represent half of the contribution for dissipated CO2 [14].

#### Development of the models

To understand the interplay between these variables, we investigated their mutual dependencies with a large number of structural equation models, as explained above. SEM is theory driven, meaning that the relationships between the variables is established on the basis of the causal relationships expected given the observed phenomena. Therefore, in a first stage of the analysis we assessed the relationships between manifest variables. For instance, it is typically assumed that higher CO2 is a consequence of a larger number number of cells, expressed as a regression CO2 $\sim$ Cells (under this notation CO2 is endogenous and Cells exogenous variables). Nevertheless, if a large amounts of CO2 per cell reflect a more active metabolism in a given environment, we may think that more cells is a consequence of better metabolic capabilities, and hence the causal relation should be expressed as a regression Cells $\sim$ CO2. This conundrum is found for almost several pairs of manifest variables, given the complexity of microbial communities. For example, the expression of any exoenzymatic activity, e.g. $\beta$–glucosidase (G), may be higher simply because there are more cells (G $\sim$ Cells), or there are more cells because the population has better capabilities for the degradation of cellulose (Cells $\sim$ G); CO2 may be a consequence of aerobic respiration (CO2 $\sim$ ATP), or more ATP is observed because aerobic respiration is happening (ATP $\sim$ CO2), etc.

As an example of exploratory modeling to elucidate these kind of relationships, in Table 5 we show an analysis of SEM models considering as variables ATP, Cells and CO2. We compared hierarchical models in which these three variables are related as A $\sim$ B $\sim$ C in all possible combinations (six models, although symmetric models are

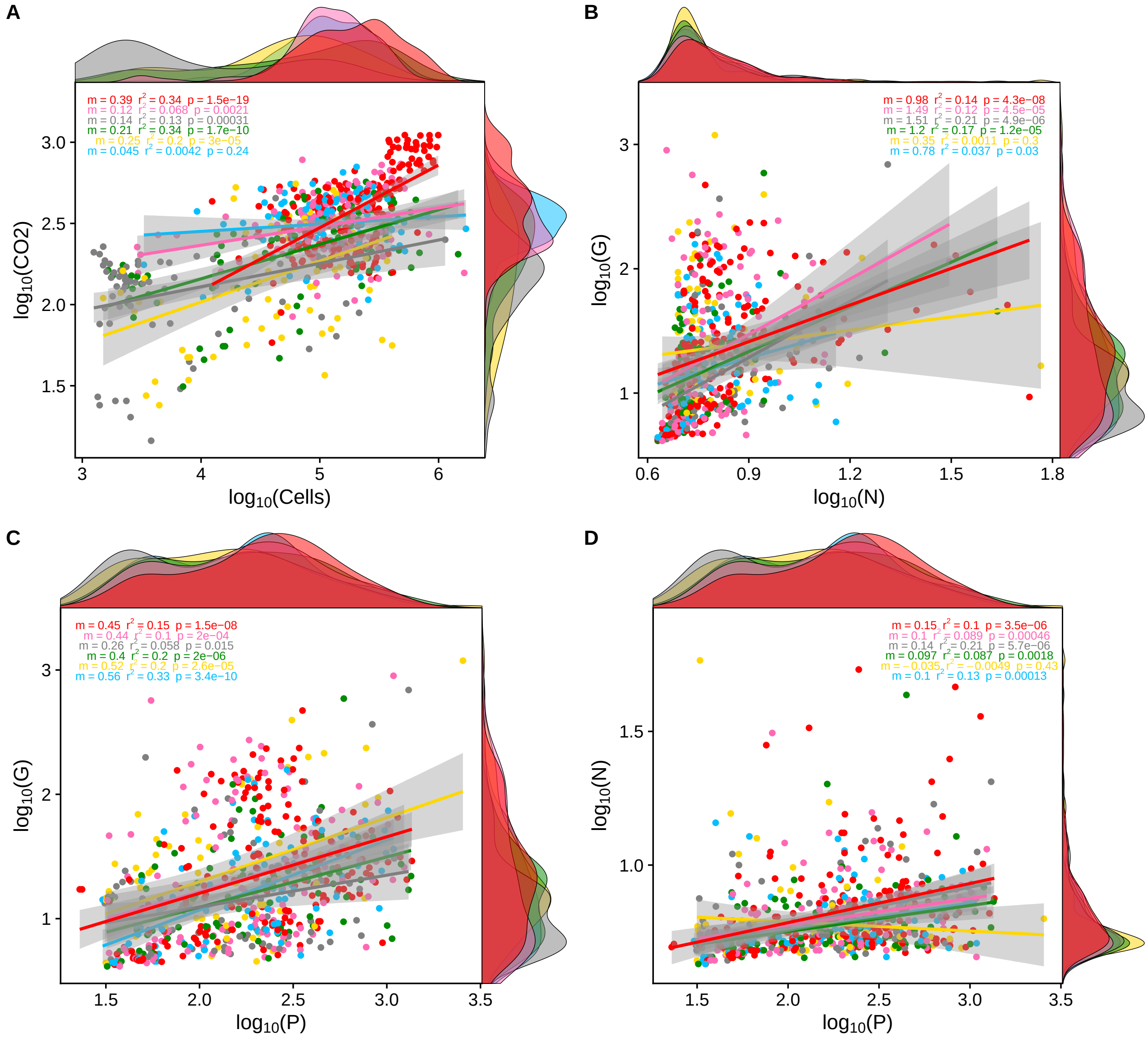


Figure 12: **Linear regressions between manifest variables.** (A) Cell counts (per million) versus $CO_2$ (ug). (B) Concentration of nitrogenase (uM) versus $\beta-$glucosidase (uM). (C) Phosphatase (uM) versus $\beta-$glucosidase (uM). (D) Phosphatase (uM) versus nitrogenase (uM). Histograms of the points differentiating the classes are located along the respective variable axes. Linear regressions are shown for each class independently, and the slope ($m$), squared correlation coefficient ($r^2$) and significance of the slope ($p$) are shown in the inset. Communities are colored according to the class they belong with the code: Class 1 = red (reference); Class 2 = green (Paenibacillus); Class 3 = pink (Klebsiella); Class 4 = yellow (Streptomyces); Class 5 = blue (Serratia); Class 6 = grey (P. putida).

| Model Id. | Structural model | Minimum test statistics | RMSEA [90% CI] |
|---|---|---|---|
| 1 | Respiration ~ Yield ~ Energy | 5.65 | 0.085 [0.028,0.16] |
| 2 | Energy ~ Yield ~ Respiration | 5.65 | 0.085 [0.028,0.16] |
| 3 | Yield ~ Respiration ~ Energy | 14.93 | 0.15 [0.087,0.22] |
| 4 | Energy ~ Respiration ~ Yield | 14.93 | 0.15 [0.087,0.22] |
| 5 | Respiration ~ Energy ~ Yield | 214.349 | 0.576 [0.51, 0.64] |
| 6 | Yield ~ Energy ~ Respiration | 214.349 | 0.576 [0.51, 0.64] |

Table 5: **Exploratory latent variable models.** Models investigating hierarchical relations between three latent variables: Energy (associated to the manifest variable ATP), Yield (cell count) and Respiration (CO2).

equivalent). Models 1 and 2 are the best choices and the only ones that are superior to the saturated counterparts, in which we also performed tests of mediation.

We also explored different definitions of latent variables, looking for manifest variables that may be simultaneously indicative of the same general process. For instance, in Suppl. Fig. 13 exoenzymatic activities were joined into the latent variable Uptake, while number of cells and amount of ATP express Yield, and CO2 Respiration. Although most of the coefficients inferred are significant the model is poor (RMSEA=0.21 with 90% CI [0.191, 0.227]). This kind of exploration led us to neglect the use of latent variables, suggesting that each measured quantity is indicative of a well differentiated processes, further confirmed by the fact that most of them took different values through the communities classes.

The best model obtained is shown in Suppl. Fig. 14 and, as we did not considered a partitioning of data into classes, it had a single coefficient for each pathway. This model represented the starting point to investigate if a model explaining better the data could be obtained under the assumption that the different classes would have different coefficients for each pathway (see Methods in Main Text and Suppl. Methods).

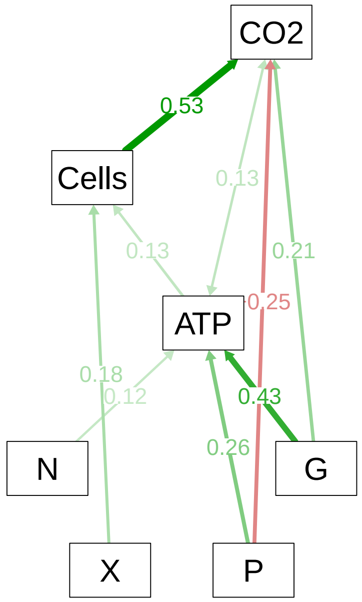


Figure 14: **Final SEM model with no classes.** Pathways and standardized coefficients for the best fit SEM obtained considering the whole dataset (i.e. without splitting into different classes). This model was then reespecified to consider classes, leading to the model presented in Fig. 3 in the Main Text.

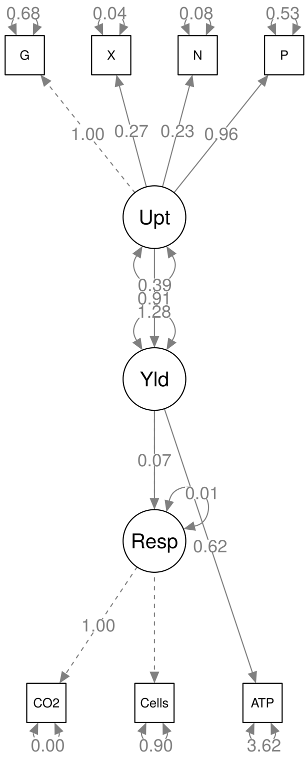


Figure 13: **Example of exploratory latent variable model.** We show one possible latent variable model in which exoenzymatic measurements are indicative of a latent variable Uptake, ATP production and cell's count of Yield and CO2 of Respiration. The structure respects the one found for the best model in Table 5, further including exoenzymatic activities. Latent variables are shown in circles, and manifest variables in squares.

| | Class 1 | | Class 2 | | Class 3 | | Class 4 | | Class 5 | | Class 6 | |
|---|---|---|---|---|---|---|---|---|---|---|---|---|
| | *Estimate* | *Z-value* | *Estimate* | *Z-value* | *Estimate* | *Z-value* | *Estimate* | *Z-value* | *Estimate* | *Z-value* | *Estimate* | *Z-value* |
| Cells~ATP | 0.128 | 4.92 | 0.128 | 4.92 | 0.128 | 4.92 | -0.024 | -0.583 | -0.024 | -0.583 | 0.128 | 4.92 |
| Cells~X | 0.915 | 3.613 | 2.304 | 5.443 | 0.184 | 0.721 | 2.304 | 5.443 | 0.915 | 3.613 | 0.915 | 3.613 |
| CO2~G | -0.028 | -7.44 | -0.028 | -7.44 | -0.028 | -7.44 | -0.065 | -7.269 | -0.028 | -7.44 | -0.028 | -7.44 |
| CO2~P | 0.016 | 2.878 | 0.016 | 2.878 | 0.016 | 2.878 | 0.039 | 8.434 | 0.039 | 8.434 | 0.039 | 8.434 |
| CO2~Cells | 0.114 | 10.168 | 0.034 | 7.239 | 0.02 | 7.525 | 0.02 | 7.525 | 0.002 | 0.204 | 0.02 | 7.525 |
| ATP~G | 0.511 | 7.998 | 0.511 | 7.998 | 0.511 | 7.998 | 0.511 | 7.998 | 0.511 | 7.998 | 0.511 | 7.998 |
| ATP~N | 0.569 | 2.719 | 0.569 | 2.719 | 0.569 | 2.719 | 0.569 | 2.719 | 1.788 | 3.27 | 1.788 | 3.27 |
| ATP~P | 0.987 | 10.577 | 1.304 | 9.948 | 0.987 | 10.577 | 0.482 | 3.31 | 1.304 | 9.948 | 0.482 | 3.31 |
| CO2~~ATP | 0.021 | 3.644 | 0.021 | 3.644 | 0.021 | 3.644 | -0.01 | -0.726 | 0.021 | 3.644 | 0.021 | 3.644 |
| Cells~~N | -0.015 | -1.06 | -0.102 | -2.506 | 0.052 | 2.362 | 0.052 | 2.362 | -0.015 | -1.06 | -0.015 | -1.06 |
| Cells~~G | -0.216 | -3.785 | -0.044 | -0.915 | -0.044 | -0.915 | -0.216 | -3.785 | -0.044 | -0.915 | -0.044 | -0.915 |
| G~~X | 0.116 | 7.948 | 0.116 | 7.948 | 0.057 | 4.007 | 0.057 | 4.007 | 0.057 | 4.007 | 0.116 | 7.948 |
| G~~N | 0.362 | 9.665 | 0.362 | 9.665 | 0.362 | 9.665 | 0.362 | 9.665 | 0.362 | 9.665 | 0.362 | 9.665 |
| G~~P | 0.073 | 6.82 | 0.134 | 8.257 | 0.196 | 5.18 | 0.073 | 6.82 | 0.073 | 6.82 | 0.134 | 8.257 |
| X~~N | 0.101 | 6.835 | 0.101 | 6.835 | 0.071 | 5.077 | -0.035 | -1.042 | 0.071 | 5.077 | 0.101 | 6.835 |
| X~~P | 0.036 | 6.241 | 0.02 | 6.462 | 0.02 | 6.462 | 0.02 | 6.462 | 0.02 | 6.462 | 0.02 | 6.462 |
| N~~P | 0.061 | 6.321 | 0.118 | 7.112 | 0.173 | 5.804 | 0.061 | 6.321 | 0.061 | 6.321 | 0.118 | 7.112 |

Table 6: **Final model coefficients**. Partial regressions are denoted as A $\sim$ B and correlations as A$\sim\sim$B. Values within the same row with the same color have the same value. Z-values $<$ abs(2.5) are highlighted in red and are not considered significant.

| Model (Parameter constraints) | | | | | | RMSEA | | | |
|---|---|---|---|---|---|---|---|---|---|
| | Estimator | df | AIC | CFI | TLI | value | 90% CI | P-val | SRMR |
| i) All equal | 294.70 | 109 | 6832.87 | 0.84 | 0.81 | 0.126 | [0.109-0.143] | <0.001 | 0.116 |
| ii) All different | 33.85 | 24 | 6742.02 | 0.99 | 0.95 | 0.062 | [0-0.107] | 0.318 | 0.026 |
| iii) Final model | 74.22 | 86 | 6658.39 | 1.00 | 1.00 | <0.001 | [0-0.035] | 0.991 | 0.051 |
| iv) Final model (shuffled) | 196.68 | 86 | 7472.67 | 0.98 | 0.97 | 0.047 | [0-0.074] | 0.540 | 0.074 |

Table 7: **Comparison of SEM models**. Summary of statistics for four SEM models considering the estimation of different coefficients for the community classes and three scenarios of parameter constraints (cases i to iii), and one additional randomization of model iii (case iv). See text for details. df=degrees of freedom; AIC=Akaike Information Criteria; CFI=Comparative Fit Index; TLI=Tucker Lewis Index; RMSEA=Root Mean Square Error of Approximation; SRMR=Standardized Root Mean Square Residual.

Respecification of the model and differentiation of the parameters by classes, led to the model shown in Fig. 3 in the Main Text, with unstandardized coefficients presented in Suppl. Table 6, highlighting with different colors those that are similar/different and their significance. As a final sanity check, we challenged the model found shuffling the samples and fitting again the model to the shuffled data. The rational is that the coefficients for the shuffled data should be similar across the groups and the model fit significantly worse than the one we found classifying the samples into their respective classes. In this way we can reaffirm the hypothesis that the different classes are truly indicative of different functional performances. In Suppl. Table 7 we provide the results found for four SEM models in which we considered the six classes and i) each coefficient is constrained to be fitted to the same value across the different classes; ii) each coefficient is completely free to be fitted to a different value; iii) an intermediate case (the final model) and iv) the final model after shuffling the samples. Model iii) stands out as the better model and, as expected, in Suppl. Fig. 7 we provide the coefficients estimated for model iv), which are approximately the same, as expected.

#### Interpretation of the SEMs

To interpret the model we considered the standardized partial regression coefficients ($m$), given that the measurements had different units. Analyzing the similarity of these coefficients led to join the classes in groups, shown in Suppl. Fig. 11. The influence of ATP on yield can be divided in two groups, with classes 1, 2, 3 and 6 depicting a significant positive relationship ($m \in [0.16 - 0.26]$, $z = 4.92$), and classes 4 and 5 with a vanishing coefficient. In

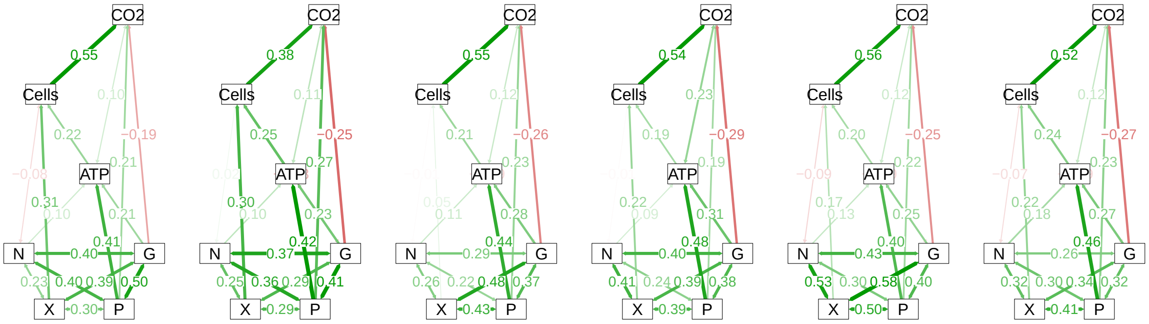


Figure 15: **Final SEM model.** Pathways and standardized coefficients for the best fit SEM obtained (shown in Fig. 3 in the Main Text) after shuffling the samples. From left to right: CLASS 1 (REFERENCE); CLASS 2 (PAENIBACILLUS); CLASS 3 (KLEBSIELLA); CLASS 4 (STREPTOMYCES); CLASS 5 (SERRATIA); CLASS 6 (P. PUTIDA).

addition, yield in class number 5 did not influence respiration, while classes 1, 2 and 6 had a significant positive relationship $m \in [0.48 - 0.59]$, $z > 7.5$. These opposite tendencies may reflect aerobic activity for classes 1, 2 and 6, while anaerobic activity would be more prevalent for class 5 (and perhaps cell death for class 6, as explained above). Indeed, increasing G simultaneously increased ATP but decreased CO2 for all classes, which may be seen as a trade-off between glycolysis and oxydative phosphorylation. The influence of exoenzymatic activities in the response variables depends on both the ability of the communities to degrade the substrate and on the amount of substrate present in the media, which was unknown. Would the substrate that the exoenzyme degrades be present in the experiments, a positive increase in the response variables with increasing exoenzymatic activity should be expected. This was the case for X, which had a positive effect on yield for classes 1, 2 and 4, with $m > 0.22$, $z > 3.6$. Xylosidase is known to be present in the extracellular matrix surrounding cellulose, and we expect it to be present in the growth media since it was made boiling fresh leaves. A positive relationship was also found between ATP and G. Nevertheless, all communities had the same unstandardized partial regression coefficient between ATP and G (see Suppl. Table 6), with modest effects across classes when the standardized coefficients were considered $m \in [0.19 - 0.32]$, $z = 7.99$. Since $\beta-$glucosidase is related with the degradation of cellulose, this observation may reflect that cellulose was present and it was abundant, and hence it did not limit growth. ATP relationship with N was significant for classes 5 and 6 with $m = 0.19$ and $m = 0.20$, $z = 3.27$ respectively. Chitinase activity is related with the presence of chitin, which is a main component of some biological structures like fungi cell walls or insect exoskeletons. Given that the substrate was only made of leaves we would not expect high levels of chitin, and therefore this relationship was noticeable because it may reflect an ecological adaptation to environments abundant in chitin, a defensive mechanism against the presence of fungi, or simply the consumption of fungus's death cells. The highly significant relationship between ATP and P was the one reflecting sharp differences between classes and a more important relative impact ($m > 0.2$ for all classes). In particular, classes 2 and 5 had the highest coefficients $m = 0.57$ and $m = 0.49$, respectively ($z = 9.95$), while classes 4 and 6 had the lowest coefficients with $m = 0.26$ and $m = 0.21$, respectively ($z = 3.31$), likely explaining the low cell count measured for these communities. The poor performance of classes 4 and 6 was likely due to a low enzymatic activity for some of their communities (see histograms in Suppl. Fig. 12), perhaps indicating an ecological adaptation to environments with a higher labile phosphorous content. This may be the case if these communities are later colonizers, due to the expected higher levels of orthophosphate [15].

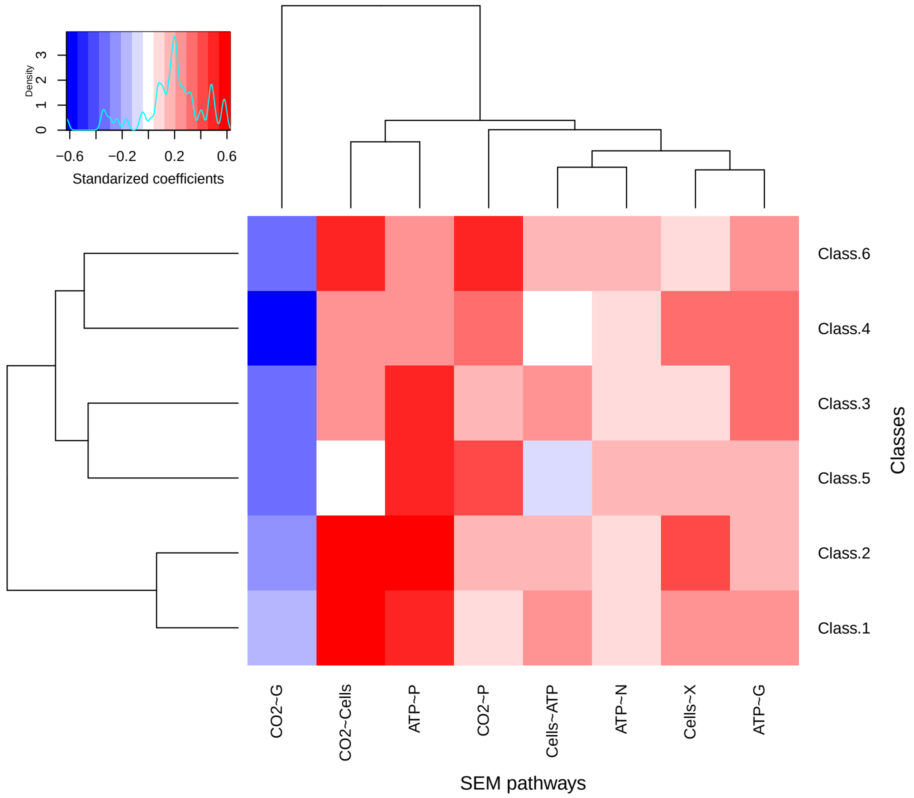


Figure 16: **Clustering of SEM standardized coefficients.** Heatmap showing the values of the standardized coefficients found for the regression pathways in the final model shown in Fig. 3 in the Main Text. Similarities between rows and columns were computed with an Euclidean distance and then clustered with the complete linkage hierarchical algorithm, implemented in HEATMAP.2 function in R package GPLOTS. No rescaling was performed.CLASS 1 (REFERENCE); CLASS 2 (PAENIBACILLUS); CLASS 3 (KLEBSIELLA); CLASS 4 (STREPTOMYCES); CLASS 5 (SERRATIA); CLASS 6 (P. PUTIDA).

| NSTI score | Low CI | Median | High CI |
|---|---|---|---|
| Class 1 | 0.060 | 0.063 | 0.067 |
| Class 2 | 0.058 | 0.062 | 0.070 |
| Class 3 | 0.056 | 0.060 | 0.064 |
| Class 4 | 0.047 | 0.051 | 0.058 |
| Class 5 | 0.045 | 0.049 | 0.055 |
| Class 6 | 0.055 | 0.061 | 0.067 |

| Num. reads | *P. borealis* | *K. pneumoniae* | *P. putida* | *S. fonticola* |
|---|---|---|---|---|
| Class 1 | 10155 | 50544 | 15476 | 34371 |
| Class 2 | 25931 | 8264 | 8123 | 22038 |
| Class 3 | 13424 | 32224 | 14042 | 24033 |
| Class 4 | 10953 | 4924 | 5825 | 4042 |
| Class 5 | 11855 | 9749 | 3779 | 80690 |
| Class 6 | 3552 | 1588 | 7790 | 2408 |

Table 8: **Additional control analysis for metagenomics predictions.** (Top) Median and lower and upper confident intervals for the NSTI scores for each class. (Bottom) Estimation of absolute cell counts (relative 16S rRNA abundances times total number of cells measured) for four abundant OTUs. The median value among the samples belonging to each class is reported (cells/ml). Class 1 (reference); Class 2 (Paenibacillus); Class 3 (Klebsiella); Class 4 (Streptomyces); Class 5 (Serratia); Class 6 (P. putida).

### 3 Metagenomic analysis

#### 3.1 Metagenomics prediction

We inferred metagenomics predictions using PiCRUST v1.1.2 [16], retrieving a large number KEGG's ortholog annotations [17]. For each sample we computed the Weighted Nearest Sequenced Taxon Index (NSTI) [16], which quantifies the average of the branch lengths that separate every OTU in the sample from the closest ortholog with a reference genome, obtaining a median among samples of 0.059, which indicates that we typically find a genome at the same genus level (95% of similarity), which we considered acceptable since it has been shown it is an ecologically coherent taxonomic level [18]. To discard potential biases arising from the different composition of the classes, we computed the median of the NSTI for each class, shown in Suppl. Table 8, top Table. We observed that most of the confidence intervals overlap, having classes 4 and 5 significantly lower values (i.e. a more reliable prediction) than the rest of the classes (Bonferroni-corrected Wilcoxon test $P < 0.01$). Importantly, we do not observed any class with a significantly worst NSTI scores than the NSTI score value observed from the whole dataset which, as we pointed out, is appropriate. In addition, since some of the metagenomic findings are obtained through relative values, we controlled that the most abundant OTUs differ in absolute values across the classes (Suppl. Table 8, right).

We observed three clear groups of predicted genes: those very abundant in every community, those that are very rare, and an intermediate group (see Suppl. Fig 17). While high frequency genes likely reflect housekeeping functions, low frequency genes are typically found in loci with high turnover rates [19] and, although they may be associated with interesting ecological processes such as the presence of phages, it is unlikely to find a robust differential statistical signal through the communities. Therefore, we focused on the group of genes found at intermediate frequencies, that are likely related with environmentally-related processes, in particular metabolic genes [19], and that our analysis confirmed.

To investigate the relationship between the genes predictions and SparCC clusters, we performed different statistical tests using STAMP [20], examining differences of the proportions of genes found in the different communities either following the KO annotations, or the second and third level defined in the KEGG pathway db. [21]. When comparisons between pairs of communities were considered we used Bonferroni-corrected p-values lower than 0.05 for Welch's tests [22][23]. Multiple communities comparisons were assessed making ANOVA tests, which we consider rejected if the Bonferroni corrected p-value was lower than 0.05, and then pairwise post-hoc tests using the Games-Howell statistics were performed, considering significant tests with p-values lower than 0.05. We further filtered out those cases were the effect sizes were lower than 0.2.

#### 3.2 PCA analysis

To get deeper mechanistic insight we performed metagenomic predictions with PiCRUST [16], that we further analyses with STAMP [20]. We selected a subset of genes such that were not very frequent nor very rare, and thus more likely representative of ecological functions (see Suppl. Methods). First of all, we performed a series of Principal Component Analysis (PCA) considering samples coming from all the classes or specific combinations of classes (see Main Text and Suppl. Fig. 18). Analysis of all classes together highlights *Pseudomonas* and *Serratia* classes as the main drivers of variability (first principal component explaining 70.4% of the variance), followed by the *Paenibacillus* and *Pseudomonas* classes again (second principal component, explaining 16.2%) of the variance, and the third component determined by the Paenibacillus class (8.5%). Therefore, the *Klebsiella* classes remain in the middle of the components and, given they represent almost half of the total communities, they may be considered representatives of the typical tree-holes communities. This brings further support to our choice of class number one as the reference class in the analysis of SEM pathways.

Pairwise PCA analysis confirms that the similarity observed in terms of composition of *Klebsiella* classes is translated into a similar genetic repertoire, as expected (see Suppl. Fig 19). For the *Pseudomonas* and *Paenibacillus* classes, the fourth class (yellow) seems to contain the largest repertoire, with more disperse points in the plot and particularly enriched in the third component (see Suppl. Fig. 20). Therefore, classes 2 (green) and 6 (gray) concentrate more genes in some of the axes determined by the yellow one. Finally, the *Serratia* class (blue) seems to be orthogonal to the gray *Pseudomonas* communities, as we will confirm in the following (see Suppl. Fig. 21).

#### 3.3 Post-hoc analysis of KEGGS's pathways

To understand the details of these trends, we clustered the KEGG's ortholog annotations into KEGG's pathways, and analyzed the results considering these classes. In Suppl. Fig. 22, we showed pairwise comparison of the classes, schematically summarized in Fig. 5 in the Main Text. The most notable finding is that we confirmed that *the Serratia* class and the (gray) *Pseudomonas* class are enriched of different genetic repertoires, that can be interpreted as different ecological strategies. We also found that communities in the *Klebsiella* class are more similar to the *Serratia* class, while communities belonging to the (yellow) *Pseudomonas* class and to the *Paenibacillus* one are more similar to the *Pseudomonas* (gray) class. Therefore, broadly speaking and consistent with the PCA analysis, we can divide the classes in two large groups, discussed in the Main Text.

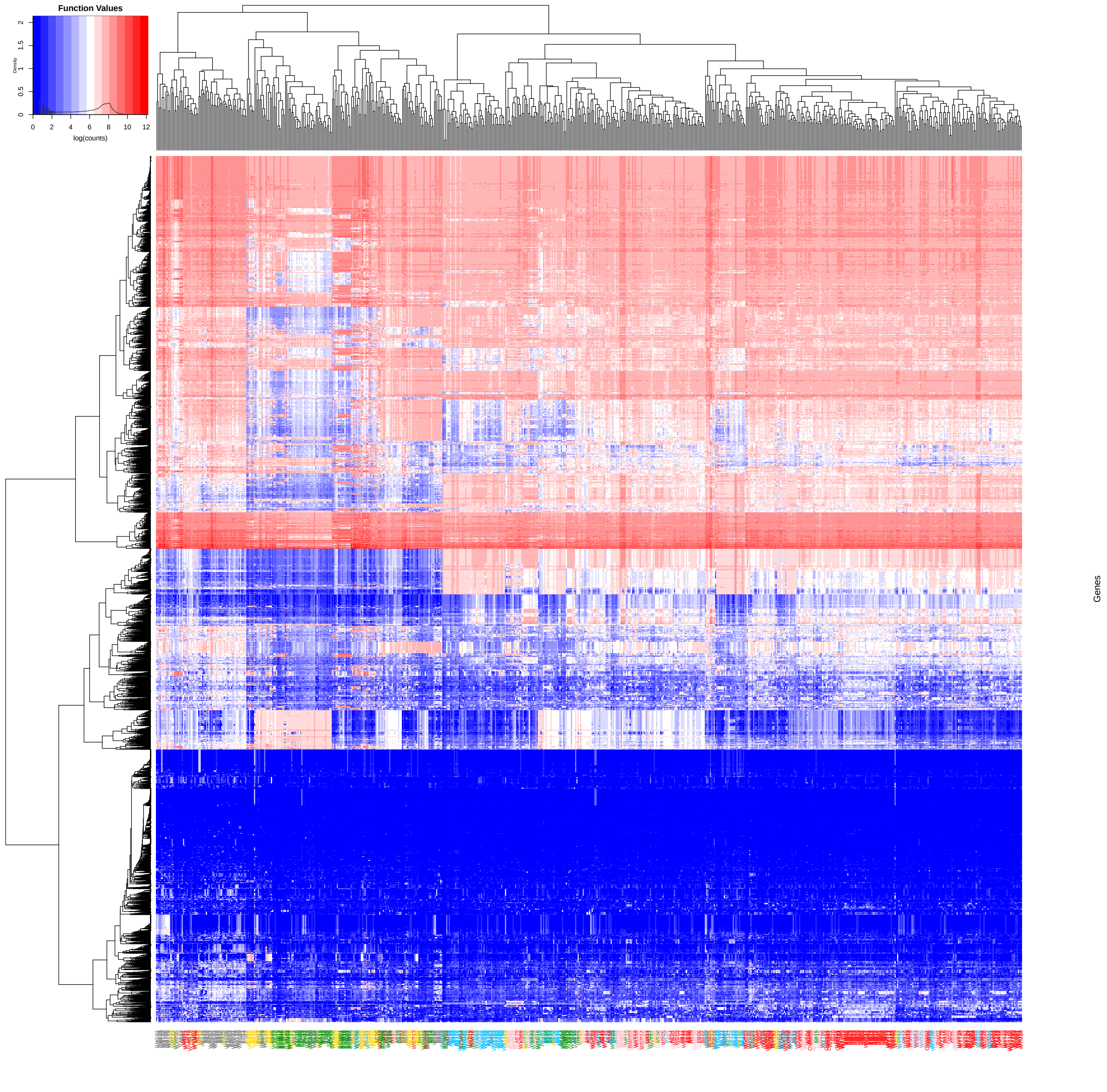


Figure 17: **Heatmap of metagenomic predictions.** Log-transformed number of gene counts (rows) for each community (columns). Clustering corresponding with the different classes is already apparent but, in order to simplify further analysis, some clusters that seem to be uniform for most of the communities were discarded (marked with red circles in the genes dendrogram). Communities are colored according to the class they belong to, with the following code: Class 1 = red (reference); Class 2 = green (Paenibacillus); Class 3 = pink (Klebsiella); Class 4 = yellow (Streptomyces); Class 5 = blue (Serratia); Class 6 = grey (P. putida).

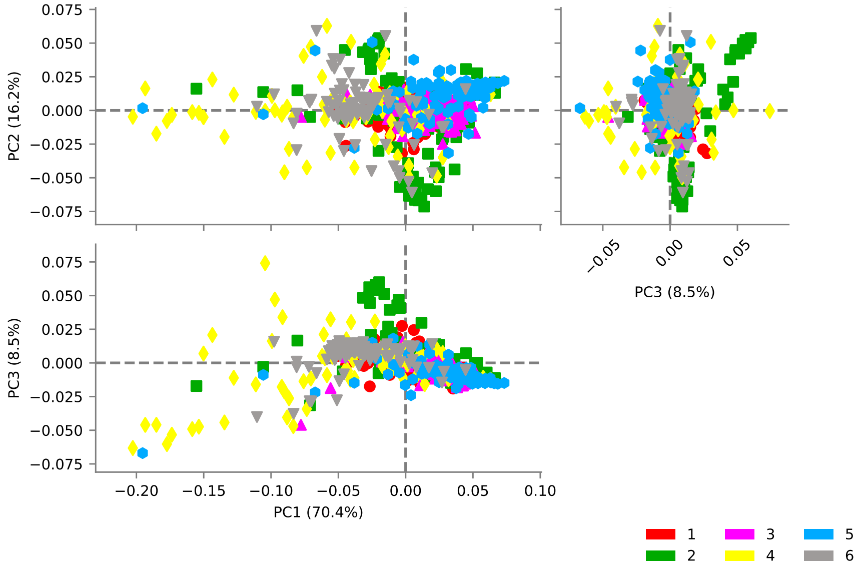


Figure 18: **Principal Component Analysis of metagenomics predictions** for all classes highlights the *Streptomyces* class (4, yellow) and the *Serratia* class (5, blue) as the main drivers of variability, spanning the first component, also contributed by class 6 (gray). The second component is mostly driven by *Paenibacillus* class (2, green) which also has points between component two and three which, together with the yellow class, are the main source of variability in that component. Class 1 = red (reference); Class 2 = green (Paenibacillus); Class 3 = pink (Klebsiella); Class 4 = yellow (Streptomyces); Class 5 = blue (Serratia); Class 6 = grey (P. putida).

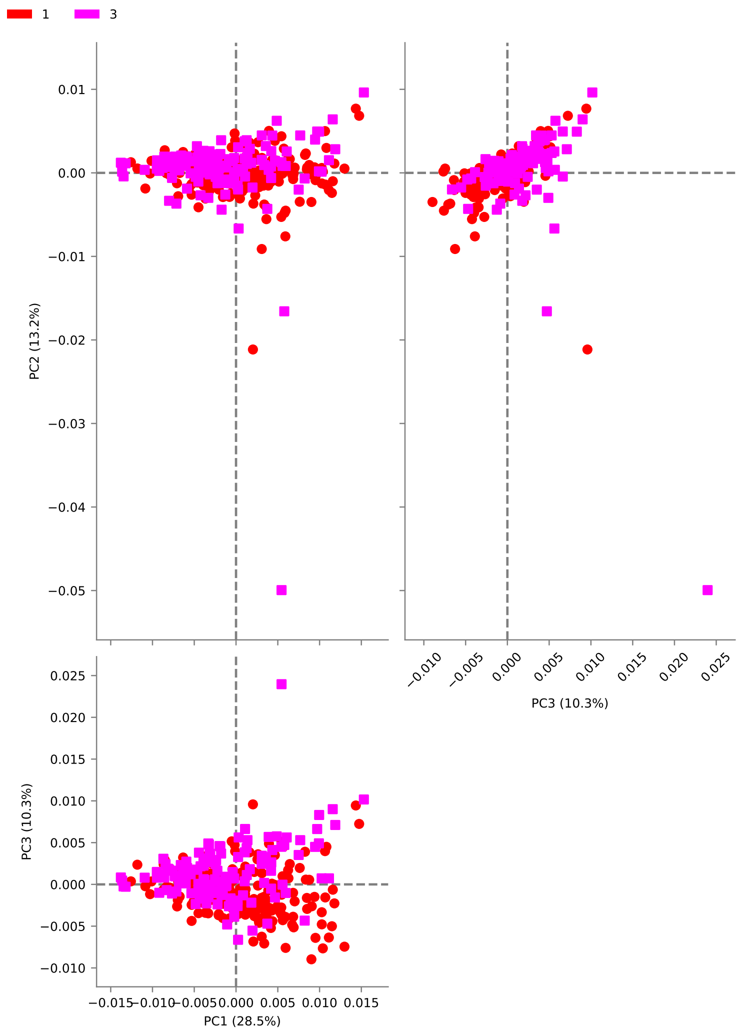


Figure 19: **Principal Component Analysis of metagenomics predictions** for classes 1 (red) and 3 (pink) confirms the similarity of both classes from the point of view of their genetic repertoire, and its centrality in the main plot. Class 1 = red (reference); Class 2 = green (Paenibacillus); Class 3 = pink (Klebsiella); Class 4 = yellow (Streptomyces); Class 5 = blue (Serratia); Class 6 = grey (P. putida).

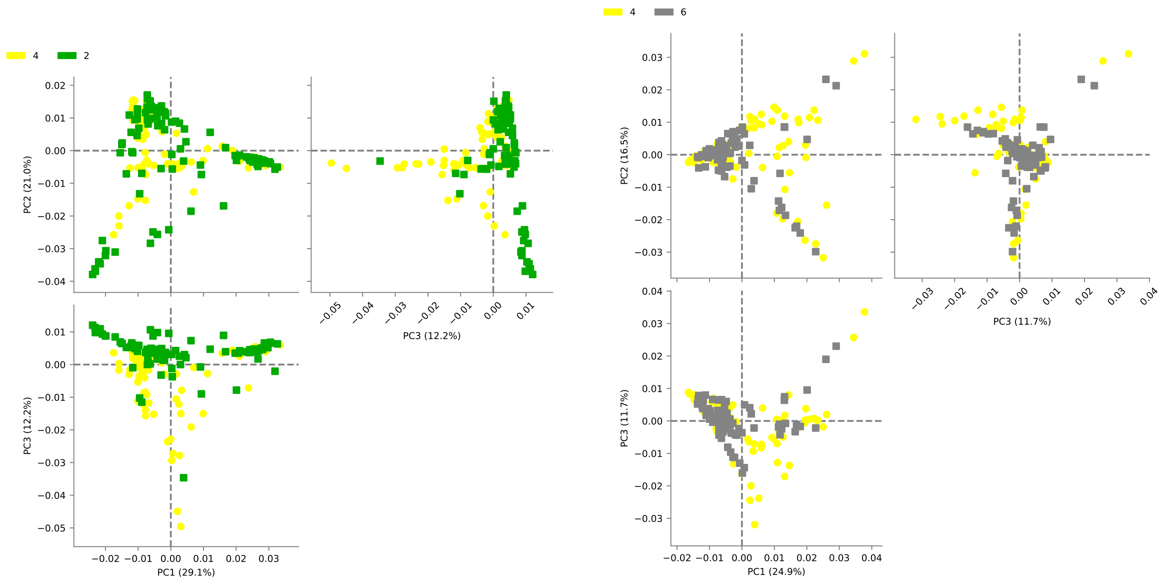


Figure 20: **Principal Component Analysis of metagenomics predictions** for classes 2 (green) and 4 (yellow) and 6 (gray). The yellow community is more central and it is used as a reference for both comparisons. Classes 2 and 6 concentrate in particular axis of the yellow community, so we expect these communities to have some degree of overlap in their genetic repertoire. Class 1 = red (reference); Class 2 = green (Paenibacillus); Class 3 = pink (Klebsiella); Class 4 = yellow (Streptomyces); Class 5 = blue (Serratia); Class 6 = grey (P. putida).

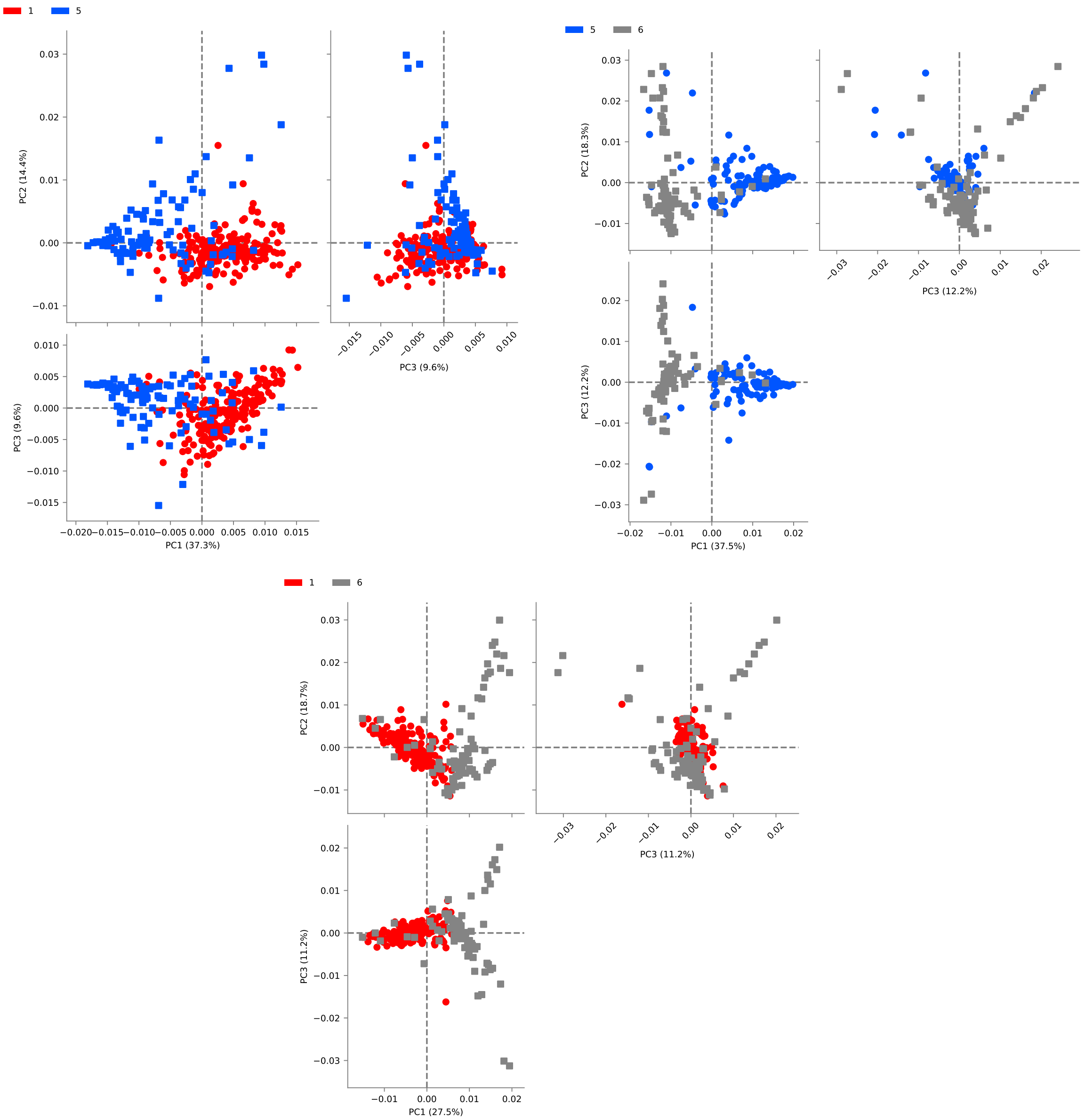


Figure 21: **Principal Component Analysis of metagenomics predictions** for classes 1 (red), 5 (blue) and 6 (gray). These three classes are well separated from each other in at least one of the components. Grey communities in particular seems to span one component were the other two are absent. Class 1 = red (reference); Class 2 = green (Paenibacillus); Class 3 = pink (Klebsiella); Class 4 = yellow (Streptomyces); Class 5 = blue (Serratia); Class 6 = grey (P. putida).

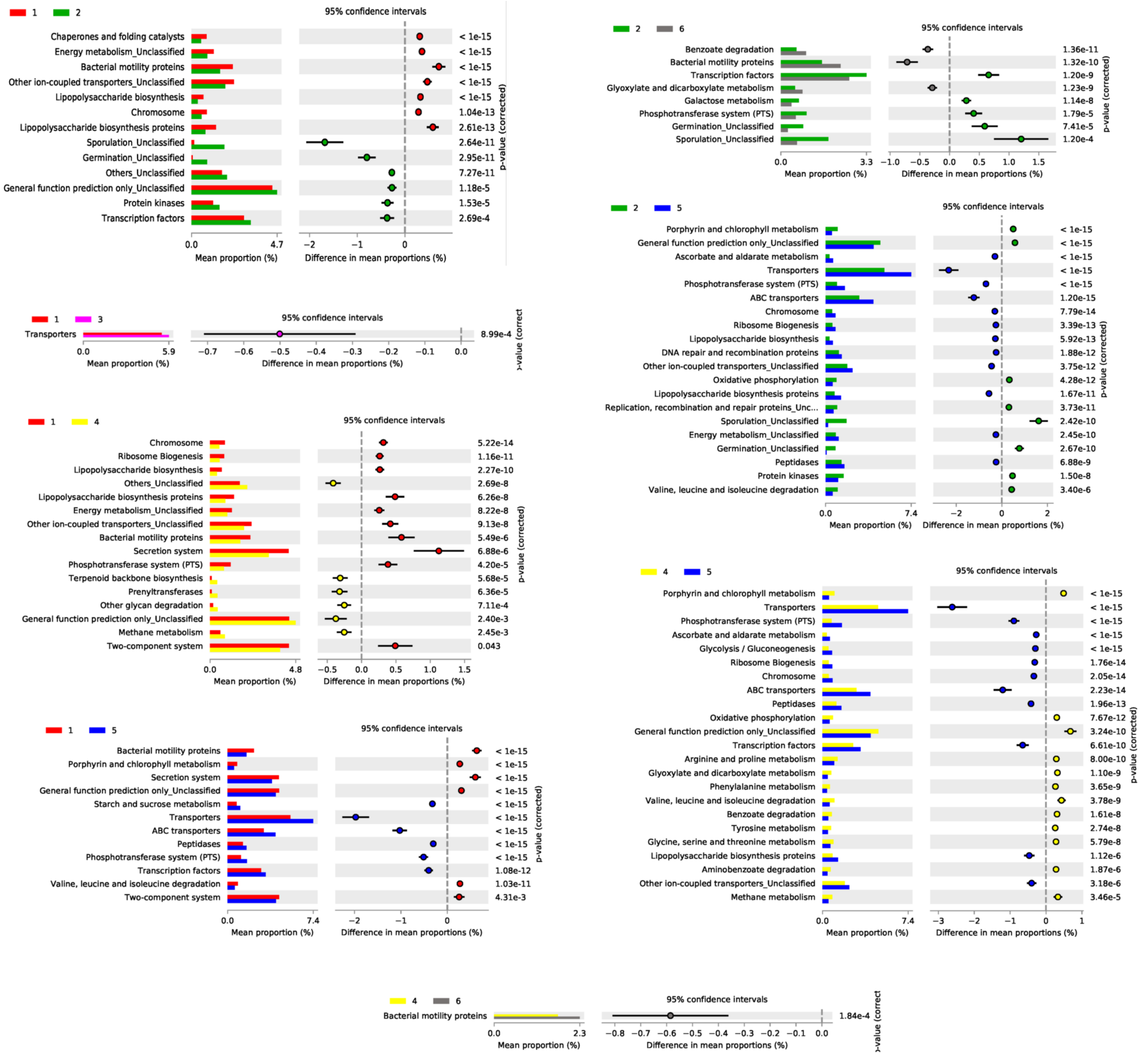


Figure 22: **Metagenomic analysis.** Comparison of the relative gene abundances of the six community classes at the third KEGG's pathway level. Each box represents the comparison of two classes, indicated by their color and number. The left column of the box indicates the mean proportion of the correspondent KEGG's pathway, and the right side the difference in mean proportions (positive if larger for the first class and negative if larger for the second one) with the correspondent corrected p-value (See Methods). As classes one and three, and classes four and six are similar, only the most informative comparisons are shown. Communities are colored according to the class they belong to, with the following code: Class 1 = red (reference); Class 2 = green (Paenibacillus); Class 3 = pink (Klebsiella); Class 4 = yellow (Streptomyces); Class 5 = blue (Serratia); Class 6 = grey (P. putida).

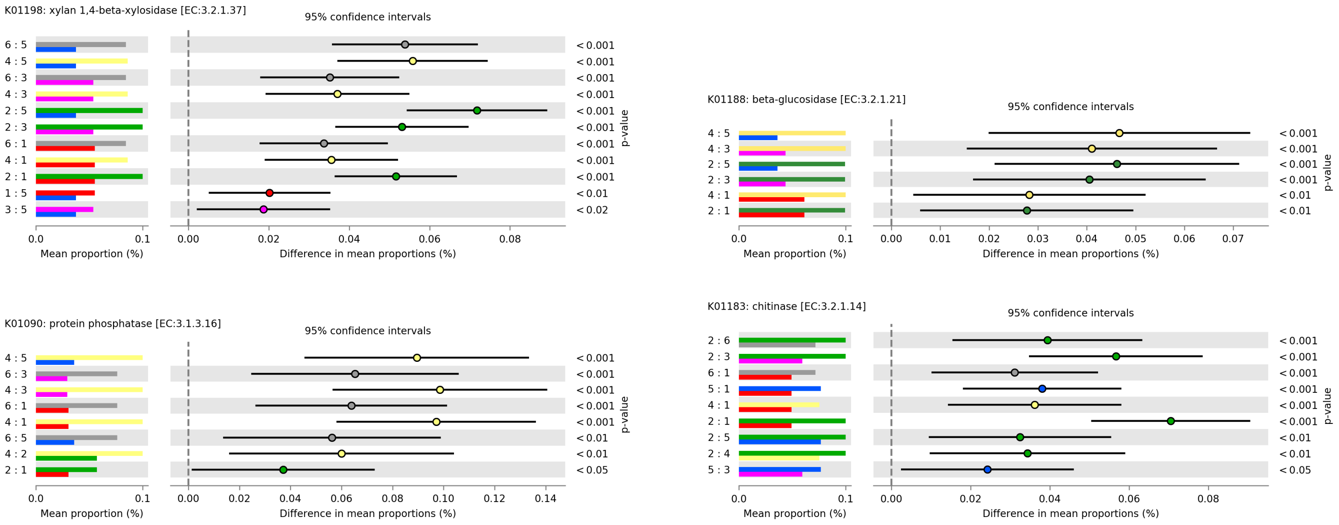


Figure 23: **Post-hoc analysis of exoenzymatic genes predictions.** The significance of the differences in mean proportions of $\beta$-xylosidase (top left), $\beta$-glucosidase (top right), phosphatase (bottom left) and chitinase (bottom right) are tested across the community classes. Each line in the figures represents a pairwise test labeled as class A:class B, and the mean proportion of each class and the difference between both are shown, together with the p-value of the test, corrected against multiple testing. Only significant comparisons (p-value$<$0.01) are shown. This analysis allow us to rank the communities for each gene or KEGG pathway according to their mean proportion. For instance, the rank for $\beta$-xylosidase would be Class 2 (green) $>$ Class 4 (yellow) $>$ Class 6 (grey) $>$ Class 1 (red) $>$ Class 3 (pink) $>$ Class 5 (blue). This figure is summarized in Fig. 4 in Main Text. Class 1 = red (reference); Class 2 = green (Paenibacillus); Class 3 = pink (Klebsiella); Class 4 = yellow (Streptomyces); Class 5 = blue (Serratia); Class 6 = grey (P. putida).

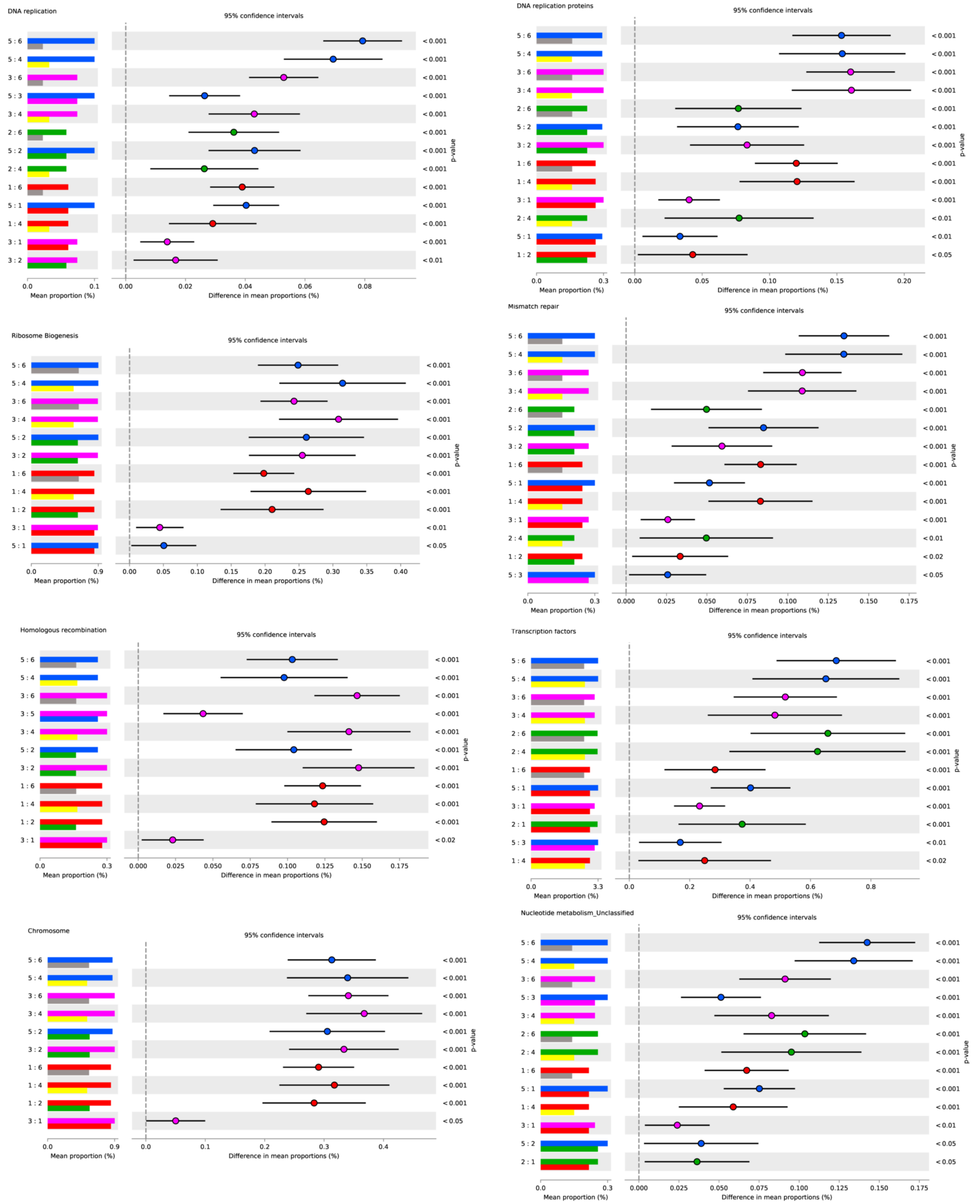


Figure 24: **Post Hoc metagenomics analysis of genetic information processing and nucleotide metabolism pathways.** The *Serratia* class followed by the *klebsiella* and *paenibacillus* classes are enriched of these annotations that likely encode capabilities for fast growth. Communities are coloured according to the class they belong with the code: Class 1 = red (reference); Class 2 = green (Paenibacillus); Class 3 = pink (Klebsiella); Class 4 = yellow (Streptomyces); Class 5 = blue (Serratia); Class 6 = grey (P. putida).

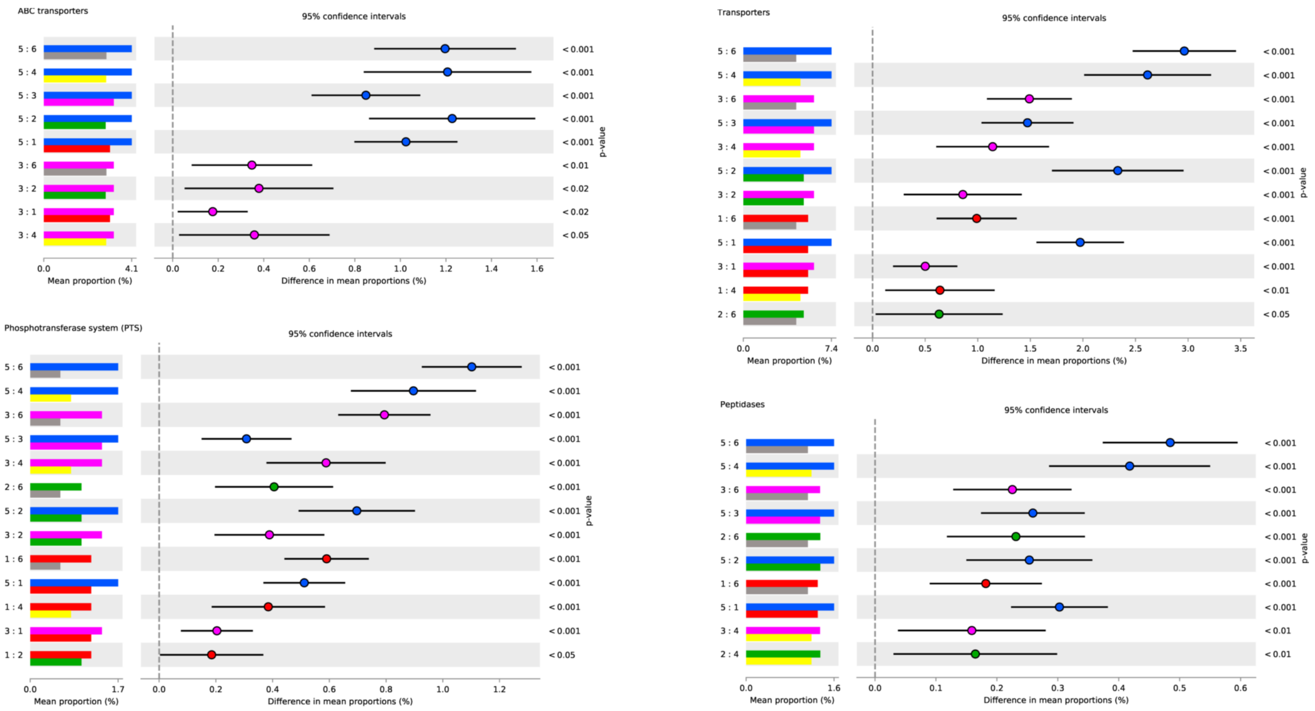


Figure 25: **Post Hoc metagenomics analysis of membrane transport and peptidases pathways.** The *Serratia* class followed by the *Klebsiella* and, in lesser extent, *Paenibacillus* classes are enriched of these annotations that likely encode capabilities for nutrient uptake from the environment. Class 1 = red (reference); Class 2 = green (Paenibacillus); Class 3 = pink (Klebsiella); Class 4 = yellow (Streptomyces); Class 5 = blue (Serratia); Class 6 = grey (P. putida).

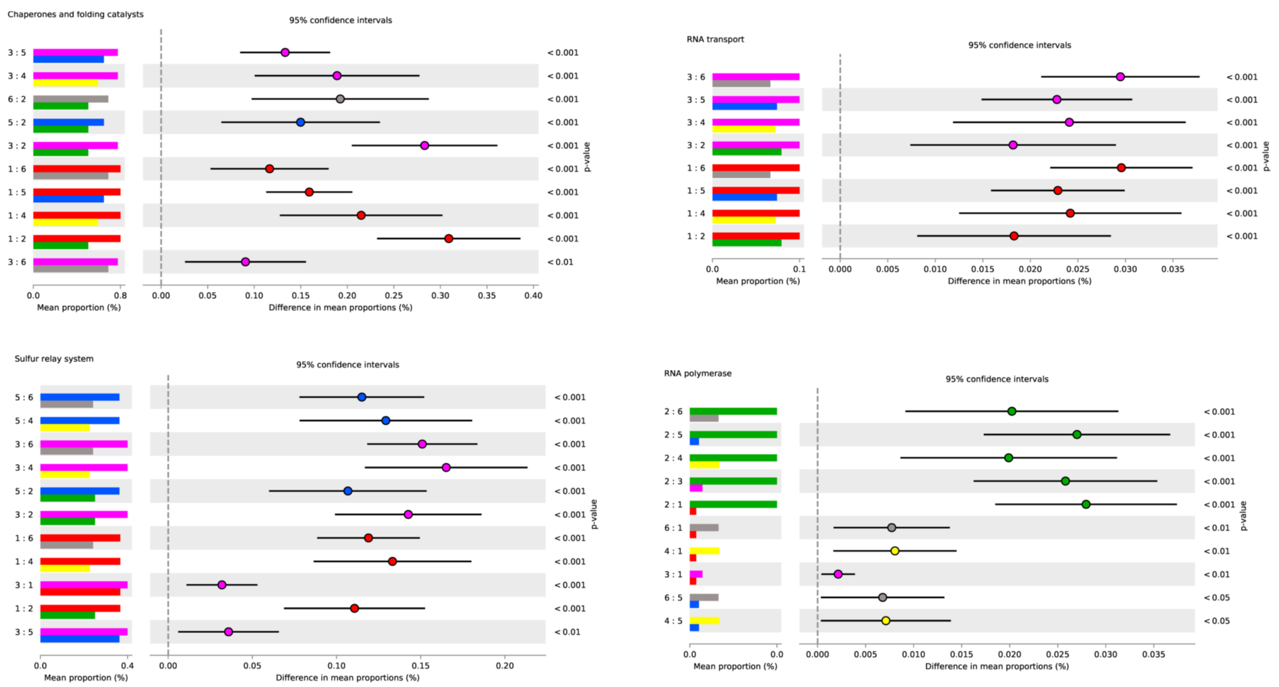


Figure 26: **Post Hoc metagenomics analysis of transcription, folding and degradation pathways.** The *Klebsiella* classes dominate these pathways, followed by the *Serratia* class, with the exception of RNA polymerase, dominated by the *Paenibacillus* community, and likely related with sporulation (see Suppl. Fig. )31. Class 1 = red (reference); Class 2 = green (Paenibacillus); Class 3 = pink (Klebsiella); Class 4 = yellow (Streptomyces); Class 5 = blue (Serratia); Class 6 = grey (P. putida).

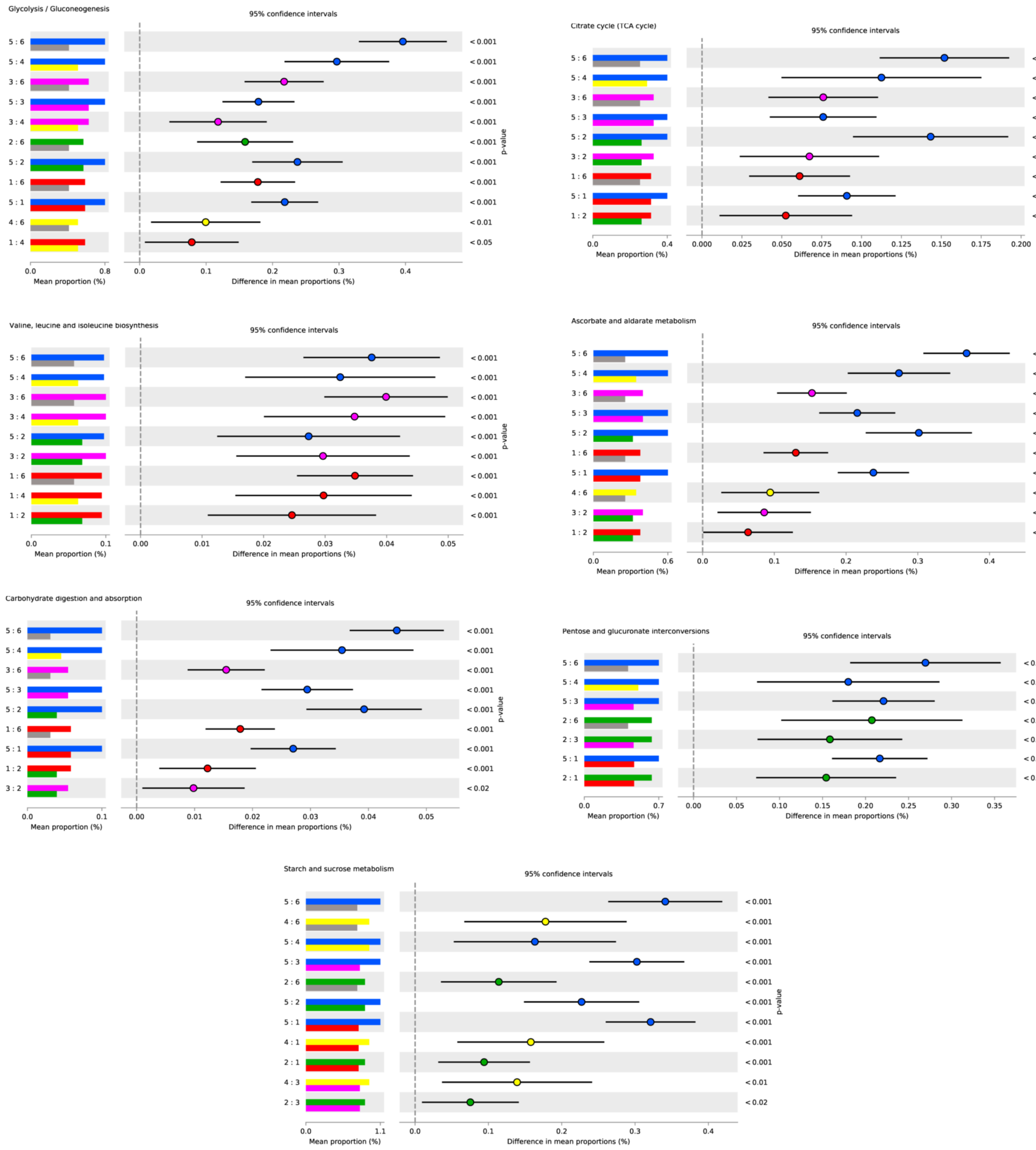


Figure 27: **Post Hoc metagenomics analysis of metabolic pathways dominated by the *Serratia* class.** The *Serratia* class followed by the *Klebsiella* classes are enriched of these annotations that encode capabilities for carbohydrate metabolism and biosynthesis of Val, Leu and Ile. Class 1 = red (reference); Class 2 = green (Paenibacillus); Class 3 = pink (Klebsiella); Class 4 = yellow (Streptomyces); Class 5 = blue (Serratia); Class 6 = grey (P. putida).

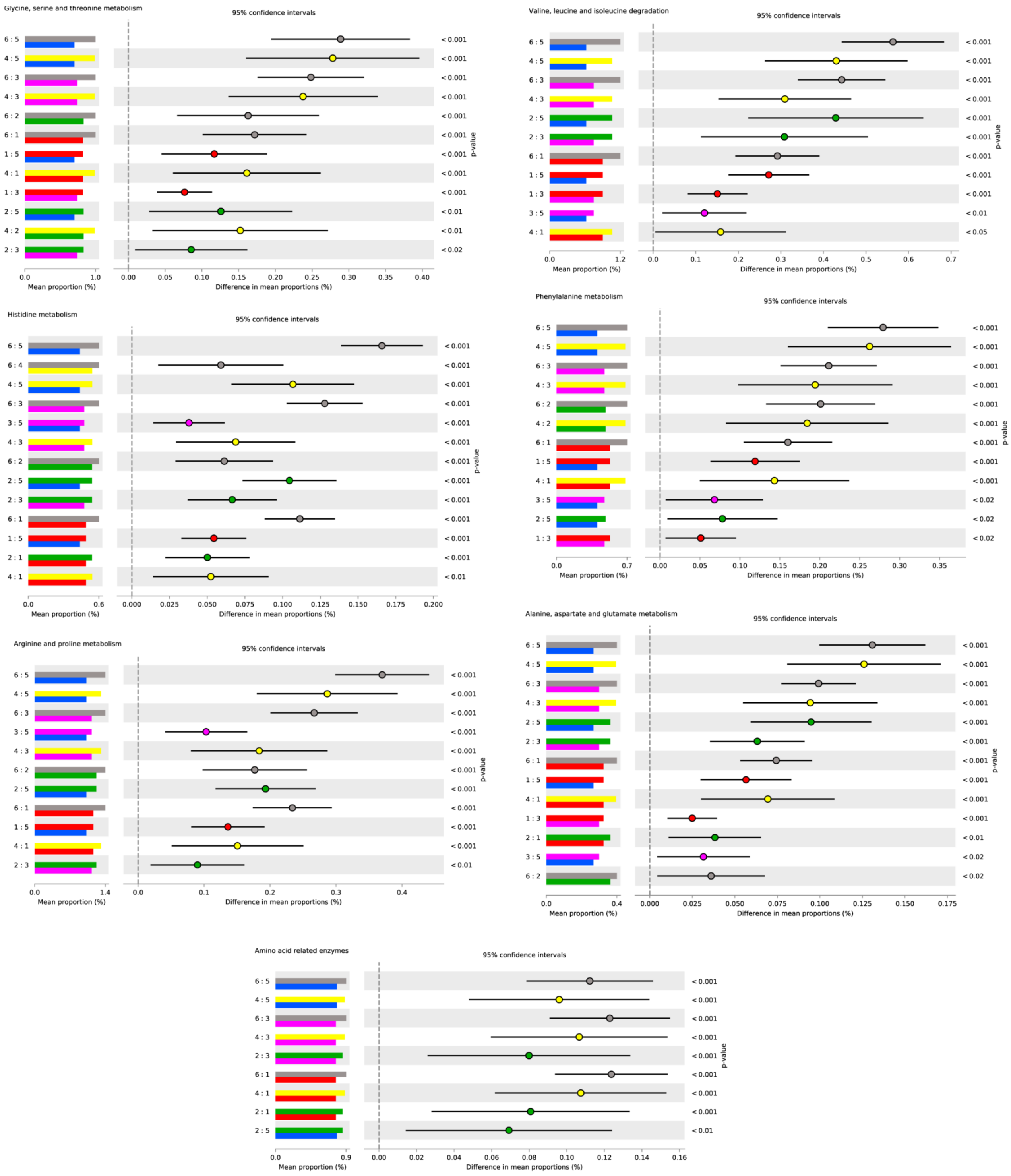


Figure 28: **Post Hoc metagenomics analysis of amino acid metabolic pathways dominated by the *Pseudomonas* (grey) class.** The *Pseudomonas* classes followed by the *Paenibacillus* and, in less extent, *Klebsiella* classes are enriched of these annotations that encode capabilities for amino acide metabolism. The *Serratia* class has in all cases the lowest number of genes of these types and, interestingly, while here we observe a large number of genes for degradation of Val, Leu and Ile, the *Serratia* has the largest pool of genes for the biosynthesis of these amino acids (see Suppl. Fig. 27). Class 1 = red (Reference); Class 2 = green (Paenibacillus); Class 3 = pink (Klebsiella); Class 4 = yellow (Streptomyces); Class 5 = blue (Serratia); Class 6 = grey (P. putida).

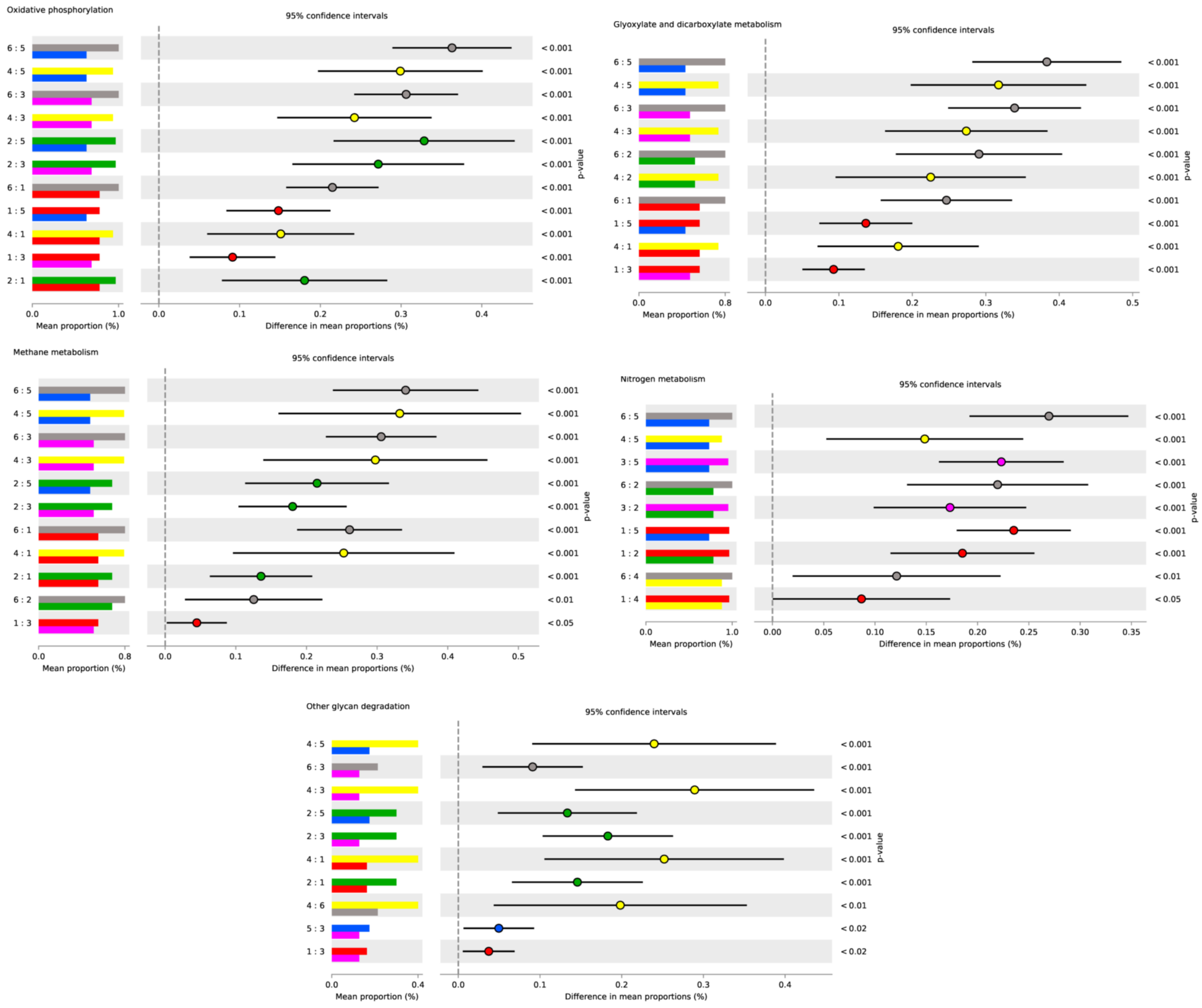


Figure 29: **Post Hoc metagenomics analysis of energy and glycan metabolic pathways dominated by *P. putida* and *Streptomyces'* classes.** The *P. putida* and *Streptomyces'* classes followed by the *Paenibacillus* and, in less extent, *Klebsiella* classes are enriched of these annotations that encode capabilities for energetic metabolism. The *Serratia* class has in all cases the lowest number of genes of these types while they have very high number of genes related with carbohydrate metabolism like glycolysis or the PPP. Class 1 = red (reference); Class 2 = green (Paenibacillus); Class 3 = pink (Klebsiella); Class 4 = yellow (Streptomyces); Class 5 = blue (Serratia); Class 6 = grey (P. putida).

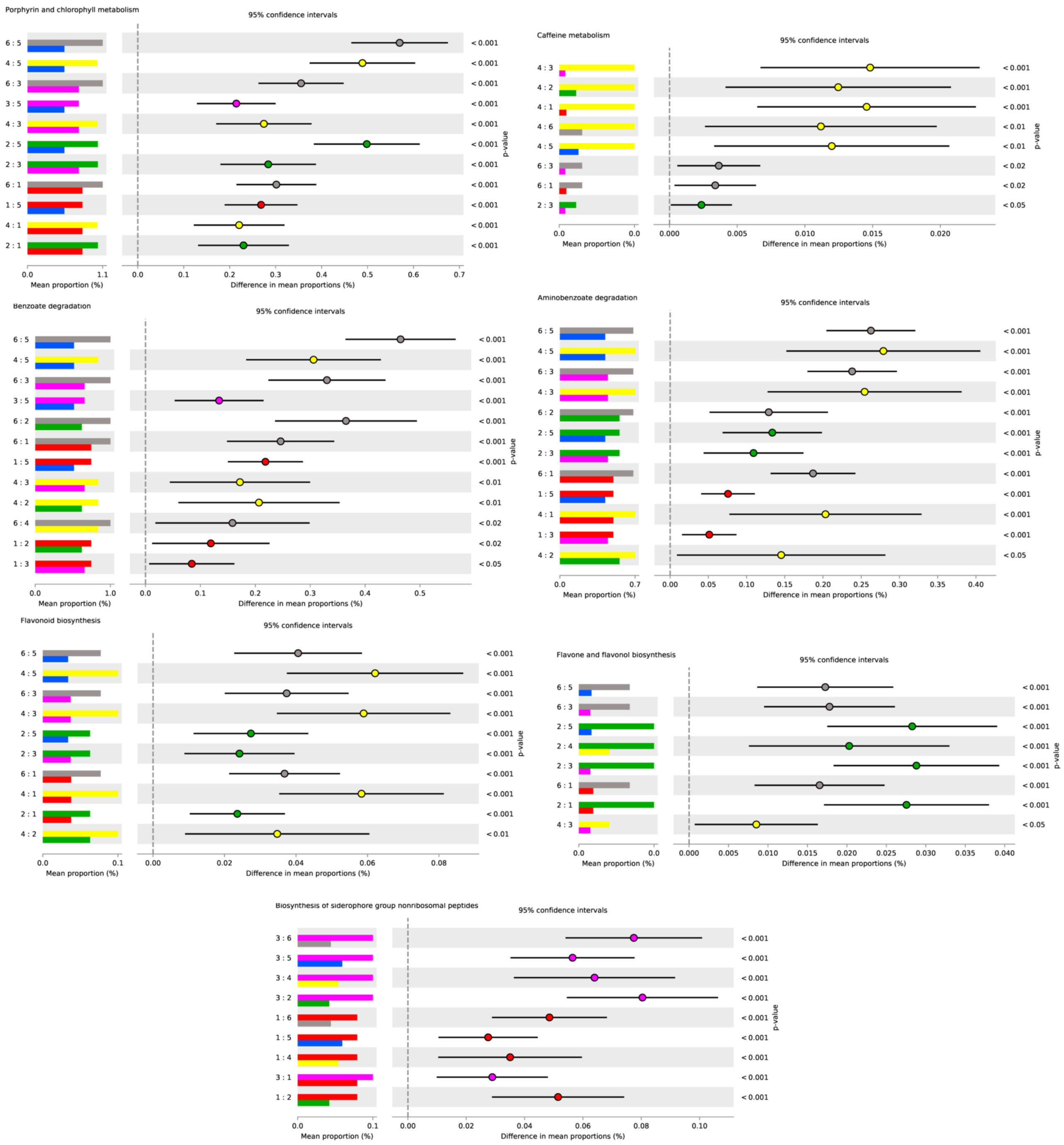


Figure 30: **Post Hoc metagenomics analysis of secondary, xenobiotic and cofactor metabolic pathways.** The *P. putida* and *Streptomyces'* classes followed by the *Paenibacillus* and, in less extent, *Klebsiella* classes are enriched of these annotations, with the notable exception of biosynthesis of siderophore group nonribosomal peptides, leaded by *Klebsiella* classes. Communities are coloured according to the class they belong with the code: CLASS 1 = RED (REFERENCE); CLASS 2 = GREEN (PAENIBACILLUS); CLASS 3 = PINK (KLEBSIELLA); CLASS 4 = YELLOW (STREPTOMYCES); CLASS 5 = BLUE (SERRATIA); CLASS 6 = GREY (P. PUTIDA).

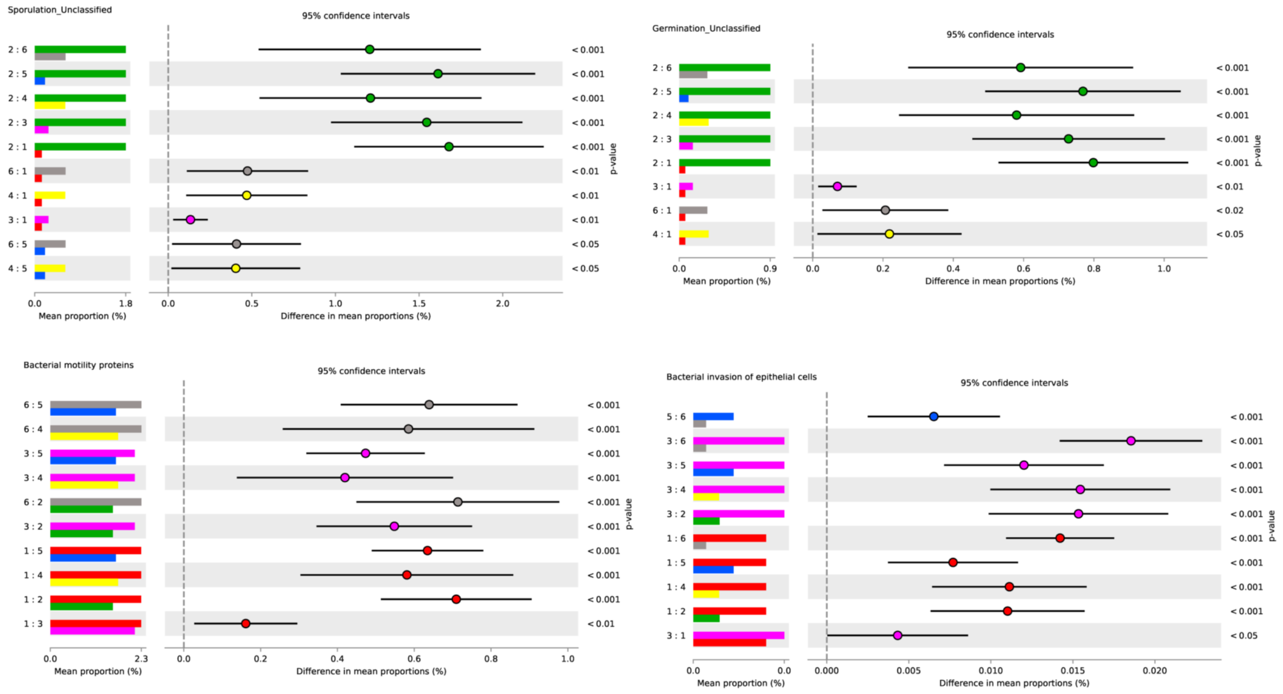


Figure 31: **Post Hoc metagenomics analysis of environmental processing and motility pathways.** The *Paenibacillus* class dominates sporulation and germination, while one of the *P. putida* and the *Klebsiella* classes lead the bacterial motility proteins pathways. The bacterial invasion of epithelial cells is related with the presence of two-component system genes and, in particular, with the ability of *Rosea* to invade organisms, which has a notable presence in *Klebsiella* classes. Communities are coloured according to the class they belong with the code: Class 1 = red (reference); Class 2 = green (Paenibacillus); Class 3 = pink (Klebsiella); Class 4 = yellow (Streptomyces); Class 5 = blue (Serratia); Class 6 = grey (P. putida).

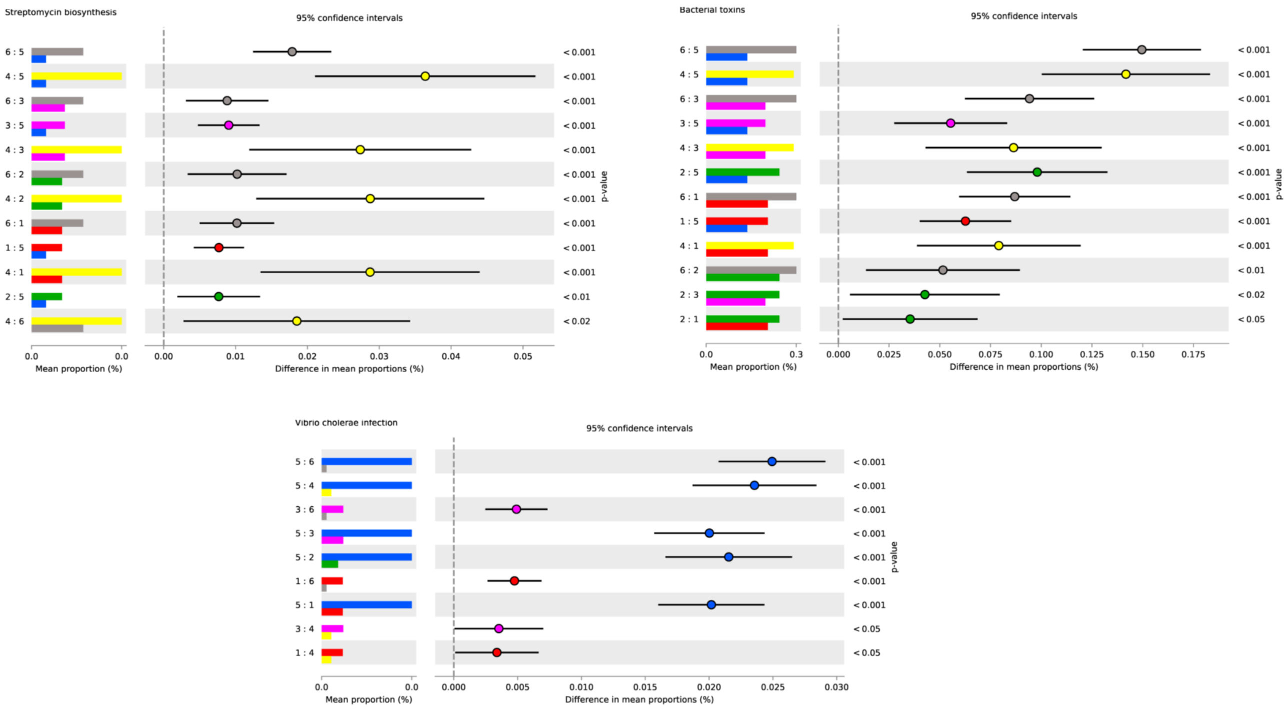


Figure 32: **Post Hoc metagenomics analysis of toxicity pathways.** The *Streptomyces* (yellow) class had a large number of genes related with streptomycin biosynthesis. The *Vibrio cholerae* infection pathway is likely explained by a large number of genes related with the type II secretion systems in the *Serratia* community. Class 1 = red (reference); Class 2 = green (Paenibacillus); Class 3 = pink (Klebsiella); Class 4 = yellow (Streptomyces); Class 5 = blue (Serratia); Class 6 = grey (P. putida).
